# Single cell RNA sequencing analysis of mouse cochlear supporting cell transcriptomes with activated ERBB2 receptor, a candidate mediator of cochlear regeneration mechanisms

**DOI:** 10.1101/2022.06.22.497230

**Authors:** Dorota Piekna-Przybylska, Daxiang Na, Jingyuan Zhang, Cameron Baker, John M. Ashton, Patricia M. White

## Abstract

Hearing loss caused by the death of cochlear hair cells might be restored through regeneration from supporting cells via dedifferentiation and proliferation, as observed in birds. We recently found that in mice, activation of ERBB2 in supporting cells promoted the differentiation of hair cell-like cells. Here we analyze transcriptomes of neonatal mouse cochlear supporting cells with activated ERBB2 using single-cell RNA sequencing. ERBB2 induction *in vivo* generated a new population of cells expressing *de novo* SIBLING (small integrin-binding ligand n-linked glycoproteins) proteins and their regulators, particularly *Secreted Phosphoprotein 1* (*SPP1*). In other systems, SIBLINGs promote cell survival, proliferation, and differentiation. ERBB2 signaling induced after noise exposure in young adult mice also up-regulated both SPP1 protein and the SPP1 receptor CD44, and drove formation of proliferating stem-like cell aggregates in the organ of Corti. Our results suggest that ectopic activation of ERBB2 signaling in cochlear supporting cells alters the microenvironment, promoting proliferation and cell rearrangements. Together these results suggest a novel mechanism for inducing stem cell-like activity in the adult mammalian cochlea.

## INTRODUCTION

Lost auditory hair cells (HCs) in adult mammals cannot be regenerated, driving permanent hearing loss. Conversely, lost HCs in birds are regenerated through the proliferation of supporting cells (SCs) and new HC differentiation (1, 2), leading to the restoration of hearing (3). Limited regenerative capacity in the organ of Corti is also observed in newborn mammals and is associated with a pool of progenitor cells in the cochlear sensory epithelium (4, 5). These include a subset of SCs in proximity to HCs, such as inner border cells (IBC), inner pillar cells (IPC), the third-row Deiter cells (DC), and lateral greater epithelial ridge (LGER) (6–8). Studies focusing on signaling pathways in mice that drive HC regeneration prior to hearing onset suggest that HC may regenerate through mitotic division followed by differentiation, via WNT signal activation (6, 9, 10). Alternatively, HCs may regenerate from progenitor cells by direct trans-differentiation, by blocking NOTCH signaling (11–14). As maturation progresses, cochlear cells become restricted from both mechanisms of regeneration (15). These findings have been interpreted to mean that even limited regeneration capacity is lost from the adult mammalian cochlea. Notably, progenitors for hair cell like cells may arise from proliferating cells in sphere cultures from adult human utricular tissue, but not adult cochlear tissues (16). Nonetheless, these findings do not rule out the possibility that quiescent adult mammalian cochlear progenitors exist (17), but require a different microenvironment to manifest their potential.

Hearing restoration in birds occurs regardless of the age. Early studies showed that signaling through phosphoinositide 3-kinase (PI3K) and the epidermal growth factor receptor (EGFR) family are required for inner ear SC proliferation in both chicken and neonatal mice (18, 19). Other studies on avian regeneration have implicated VEGF activation, which also signals through PI3K (20, 21). In the EGFR family, there are four closely related receptor tyrosine kinases: ERBB1, ERBB2, ERBB3, and ERBB4. These form homo or heterodimers with each other upon binding of the ligand and inducing molecular pathways supporting cells proliferation, migration and survival. We previously showed that signaling mediated by ERBB2, which is the preferred heterodimerization partner for the other three EGFR family members, drives proliferation of SCs in explant cultures and supernumerary hair cell-like formation *in vivo* in the neonatal mouse cochlea (22). Notably, in those studies, lineage tracing revealed that cells expressing constitutively active (CA) ERBB2 mutant induced these activities in neighboring cells, suggesting the presence of an amplifying signal cascade that recruits adjacent tissue.

Consistent with this interpretation, SOX2 protein was broadly down-regulated in the apical turn harboring sparse CA-ERBB2+ cells (22).

To identify ERBB2-mediated signaling pathways associated with the changes in SCs that may facilitate HC regeneration, we performed single cell RNA sequencing (scRNA-seq) on sorted neonatal SCs with and without CA-ERBB2. We performed cluster-specific gene analysis to identify differentially expressed genes (DEGs) in different cell populations. Our goals were to identify clusters of CA-ERBB2 cells that are transcriptionally distinct, assign clusters and DEGs to SC subtypes and identify candidate secondary signaling pathways that could act downstream of ERBB2 signaling to promote local changes. We further sought to confirm if selected downstream mediators were also expressed in the cochlea of young adult mice after damage and CA-ERBB2 activation, and if so, where they were located.

## RESULTS

### Experimental design to determine transcriptome of cochlear SCs expressing ERBB2

To perform scRNA-seq analysis of SCs expressing ERBB2, we employed the same genetic mouse model that revealed an indirect effect of ERBB2 activation on the formation of ectopic hair cell-like cells (S1 Fig) (22). These mice harbor an inducible mutated *ErbB2* transgene encoding CA-ERBB2. In order to limit activation of CA-ERBB2 to SCs, we used an inducible DNA Cre recombinase under control of the SC gene promoter *Fgfr3* (S2 Fig) (23). Double heterozygotic mice (“Tet-On” *CA-ErbB2, Fgfr3-iCre*) were then bred to homozygous line harboring a “floxed” TA transcription factor, with an IRES-GFP to use as a lineage marker (“ROSA-rtTA-GFP”). The resulting pups with *CA-ErbB2* and *Fgfr3-iCre* (denoted here as “CA-ERBB2”) were used to analyze the transcriptome of SCs with induced ERBB2 signaling; whereas pups harboring *Fgfr3-iCre* were used as “Control”. Activation of iCRE was performed by two injections of tamoxifen at P0 and P1, which resulted in GFP expression to label SCs (S1 Fig). CA-ERBB2 expression was induced at P2 by doxycycline injection. The organs of Corti were dissected from P3 pups, and living GFP+ SCs were purified by fluorescence-activated cell sorting (FACS) by excluding cells that had taken up a cell-impermeant dye (S2 Fig). On average, we obtained 55±13 GFP+ cells per cochleae. RNA cDNA libraries were prepared and sequenced from about 300 GFP+ single cells per genotype collected in three independent FACS experiments with 85% cells having less than 20% mitochondrial transcript expression (S3 Fig).

### New populations of transcriptionally distinct cochlear cells arise in response to ERBB2 signaling

Quality control metrics revealed similar numbers of transcripts and UMI counts for the two genotypes (Fig 1A). A preliminary UMAP plot of all of the cells revealed intermingling of CA-ERBB2 and Control cells as well as evidence for their separate differentiation (Fig 1A, right). Unbiased clustering was then used to group all GFP+ cells prior to assessment of transcriptome changes associated with CA-ERBB2 expression. Based on the pattern of gene expression, FACS-purified GFP+ cells can be grouped into at least 5 clusters (Fig 1B, C). Two clusters showed significant differences between Control and CA-ERBB2 samples in distribution of cochlear cells: cluster C4, mostly comprised of Control cells, and cluster C3 formed entirely by CA-ERBB2 cells. Significant changes in the cell distribution were also observed within cluster C2, where CA-ERBB2 cells have clearly segregated away from Control cells. There were no differences between Control and CA-ERBB2 in distribution of cochlear cells in clusters C0 and C1. These results suggest heterogeneity in the responses of SCs to intrinsic ERBB2 activation. In a minority of GFP+ cells, ERBB2 activation drove a greater transcriptional response, one that differentiated the activated cells away from the parent population.

**Fig 1.**
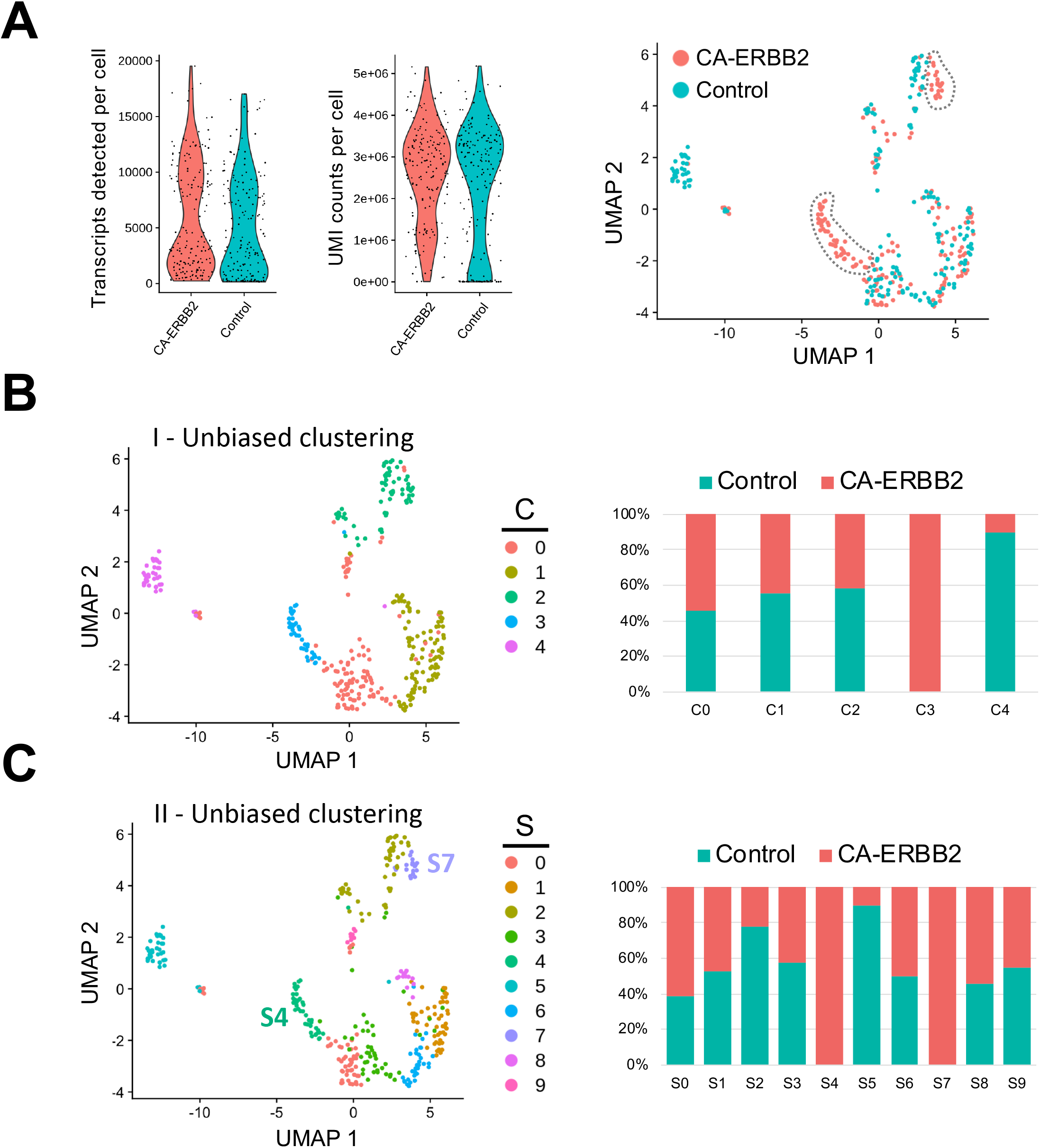
Unbiased Seurat clustering of cochlear SC transcriptomes from Control and CA-ERBB2 samples. (A) Violin plots showing similar distributions observed for cells from Control and CA-ERBB2 samples in the number of detected transcripts per cell and in UMI counts per cell used in generating sequencing libraries. On the right, preliminary UMAP plot showing distribution of CA-ERBB2 cells and Control cells. Populations of CA-ERBB2 cells that shifted away from Control cells are depicted by dotted line. (B) Initial unbiased clustering showing differences between Control cells and CA-ERBB2 cells in the number of clusters and in distribution of cells within the clusters. On right, bar graph showing proportion of Control and CA-ERBB2 cells found in each cluster. (C) A second unbiased clustering distinguishes subpopulations within clusters C0, C1 and C2, generating ten unique clusters (S0-S9) with two clusters formed by CA-ERBB2 cells only (S4 and S7) and two clusters mostly composed of Control cells (S2 and S5). On right, bar graph showing proportion of Control and CA-ERBB2 cells found in each cluster. Statistical analyses were performed with Seurat package in R version 4.1.2.

To better understand the transcriptional diversity of cochlear SC with induced ERBB2 signaling, we performed unbiased clustering that resulted in distinguishing subpopulations within clusters C0, C1 and C2. Ten unique clusters (S0-S9) were generated with two clusters composed of CA-ERBB2 cells only (Fig 1C). Notably, in addition to cluster S4 that corresponds to C3 in first unbiased clustering, CA-ERBB2 cells in cluster C2 were distinguished as cluster S7. Most of remaining cells in cluster C2 were Control cells (~80%) and were identified as cluster S2. Another cluster represented mostly by Control cells (~90%) is S5, which corresponds to C4 in the first unbiased clustering. In remaining clusters, distribution of Control and CA-ERBB2 cochlear cells was similar.

### Assignment of SC subtypes to clusters

To assess which SCs contributed to which clusters, we graphed expression of genes specific to SC subtypes, as described for scRNA-seq of P1 SCs (24) (S4 Table, S5 Fig). At birth, *Fgfr3-iCre* can be expressed in Deiter cells (DC), inner and outer pillar cells (IPC, OPC), outer hair cells (OHC), and Hensen cells (HeC). In our system, *Fgfr3-iCre* expression is sparser, and expression in HC, DC, and HeC is biased towards the apical region, where cells are less differentiated (S2 Fig). Here we used the 5-cluster analysis, as we anticipated a similar number of cell types. Results in Fig 2 show that the HC markers *Ccer2, Pvalb*, and *Insm1* are clustered together in a subset of C2 (Fig 2B, C). The HeC marker *Fst* is expressed in C1, whereas *Nupr1* is present in both C1 & C2 (Fig 2B, D). All five DC markers, *Hes5, Pdzk1ip1, S100a1, Prss23*, and *Lfng*, were most strongly up-regulated in C2 (Fig 2B, E). PC markers were more ambiguous, but were most up-regulated also in C2 (Fig 2B, F). C4, which is comprised primarily of Control cells, expressed all markers examined at a moderate level and thus could not be identified. C0 is a mix of CA-ERBB2 and Control cells, and it expresses a handful of sensory-specific markers, which were *S100a1, S100b*, and *Cryab*. Its identity is likewise ambiguous. C3 is composed exclusively of CA-ERBB2 cells and primarily expresses OPC markers (Fig 2B, F, note *Smagp* levels). However, it also contains identifying markers for LGER1 (LGER group 1), such as *Dcn, Ddost, Pdia6, Rcn3*, and *Sdf2l1*. These are also enriched in C1 and C2 (Fig 2G).

**Fig 2.**
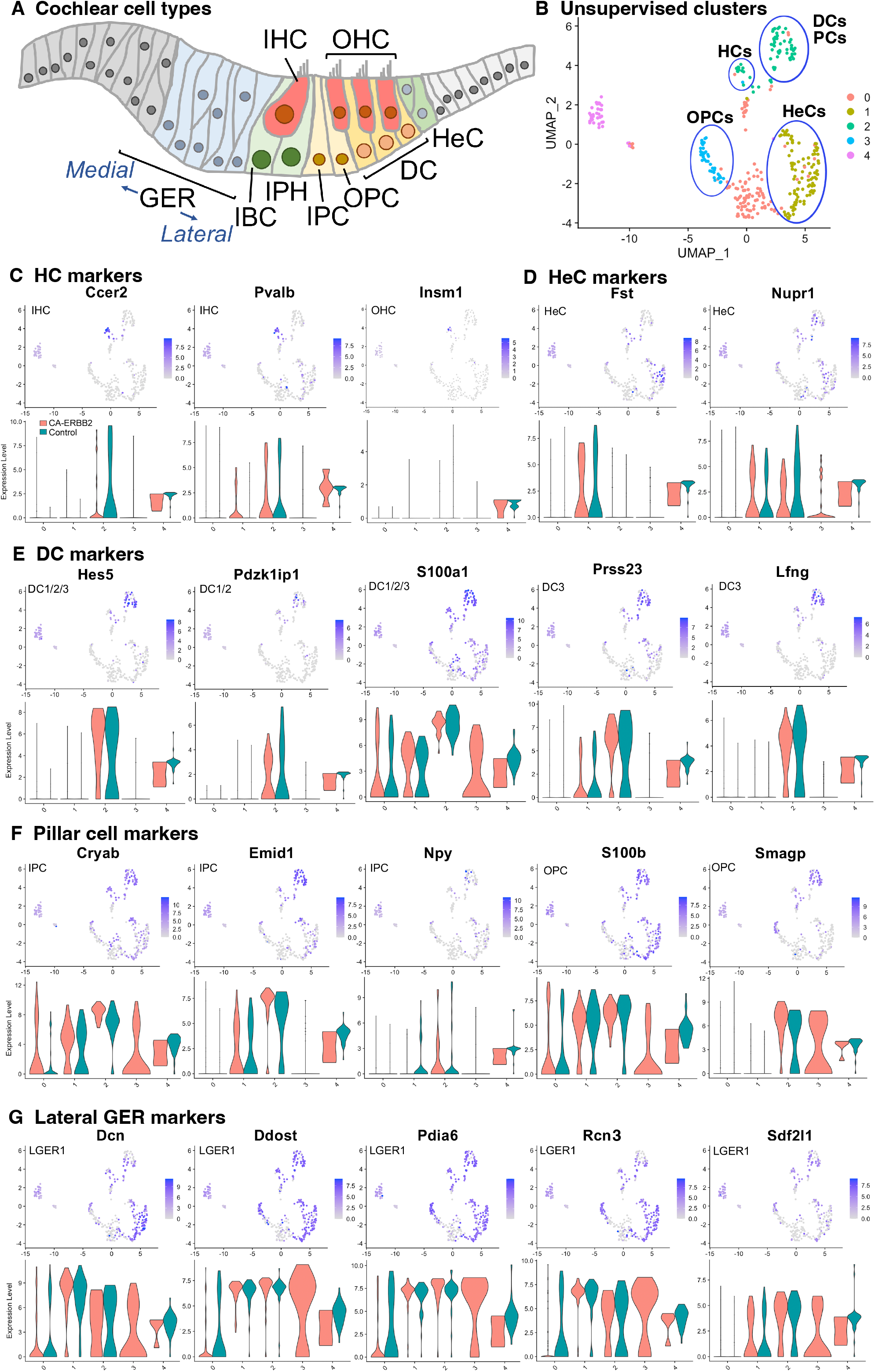
UMAP plots and violin plots for marker gene transcripts identifying cochlear cells. (A) Schematic cross-section of the cochlear duct showing the position of different cell types. IHC, inner hair cells; OHC, outer hair cells; GER, greater epithelial ridge; IBC, inner border cells; IPH, interphalangeal cells; IPC, inner pillar cells; OPC, outer pillar cells; DC, Deiter cells; HeC, Hensen cells. (B) Cluster plot from Fig 1b, with putative cell type designations. (C) Three HC markers are presented: *Ccer2, Pvalb*, and *Insm1*. (D) Two HeC markers are shown: *Fst* and *Nupr1*. (E) The following marker genes were used to identify Deiter cells: *Hes5, Pdzk1ip1, S100a1, Prss23, Lfng*. (F) Five markers for IPC and OPC are shown: *Cryab, Emid1, Npy, S100b*, and *Smagp*. (G) Lateral GER cells express *Dcn, Ddost, Pdia6, Rcn3, Sdf2l1*. The most informative markers were chosen for display. Cochlear supporting cell identity associated with analyzed marker gene is indicated on UMAP plot. Expression of the HeC and OPC marker gene *Fam159b*, and the IHC marker gene *Acbd7* were not detected.

### CA-ERBB2 SCs are transcriptionally distinct from Control SCs

Induction of ERBB2 signaling in CA-ERBB2 cells may result in activation of genes that normally are not expressed in cochlear SCs. Therefore, to identify DEGs between Control and CA-ERBB2 samples, we performed statistical analysis using both the Wilcoxon rank-sum test and the Likelihood-Ratio (LR) test. In the Wilcoxon rank-sum test, DEGs are identified among genes that have some level of expression in the compared conditions. The LR test allowed for detection of DEGs which expression is undetectable within one condition (25). Genes were mapped in the Gene Ontology resource (S6 Fig). Sequences that could not be mapped to a corresponding protein record in the PANTHER classification system were designated “unmapped IDs.” They mostly represent non-coding RNAs and pseudogenes.

Using Wilcoxon rank-sum test (adj *p* < 0.05, logFC > 2), 584 DEGs were identified, with 68 genes up-regulated and 516 genes down-regulated (S7 Data). Around half of the genes in each group were unmapped IDs. Almost 90% of DEGs were found in less than 10% of the CA-ERBB2 population. Using the LR test (adj *p* < 0.05, logFC > 2), 1556 DEGs were identified (S8 Data). 1049 genes were up-regulated, and 507 genes were down-regulated, and around half of them were protein coding genes (S6 Fig). About 64% of the DEGs were found in less than 10% of CA-ERBB2 population. Volcano plots were used to show DEGs for both analyses (Fig 3A). Note how the LR analysis enables the identification of very significant DEGs like *Wnt10a* (Fig 3A right).

**Fig 3.**
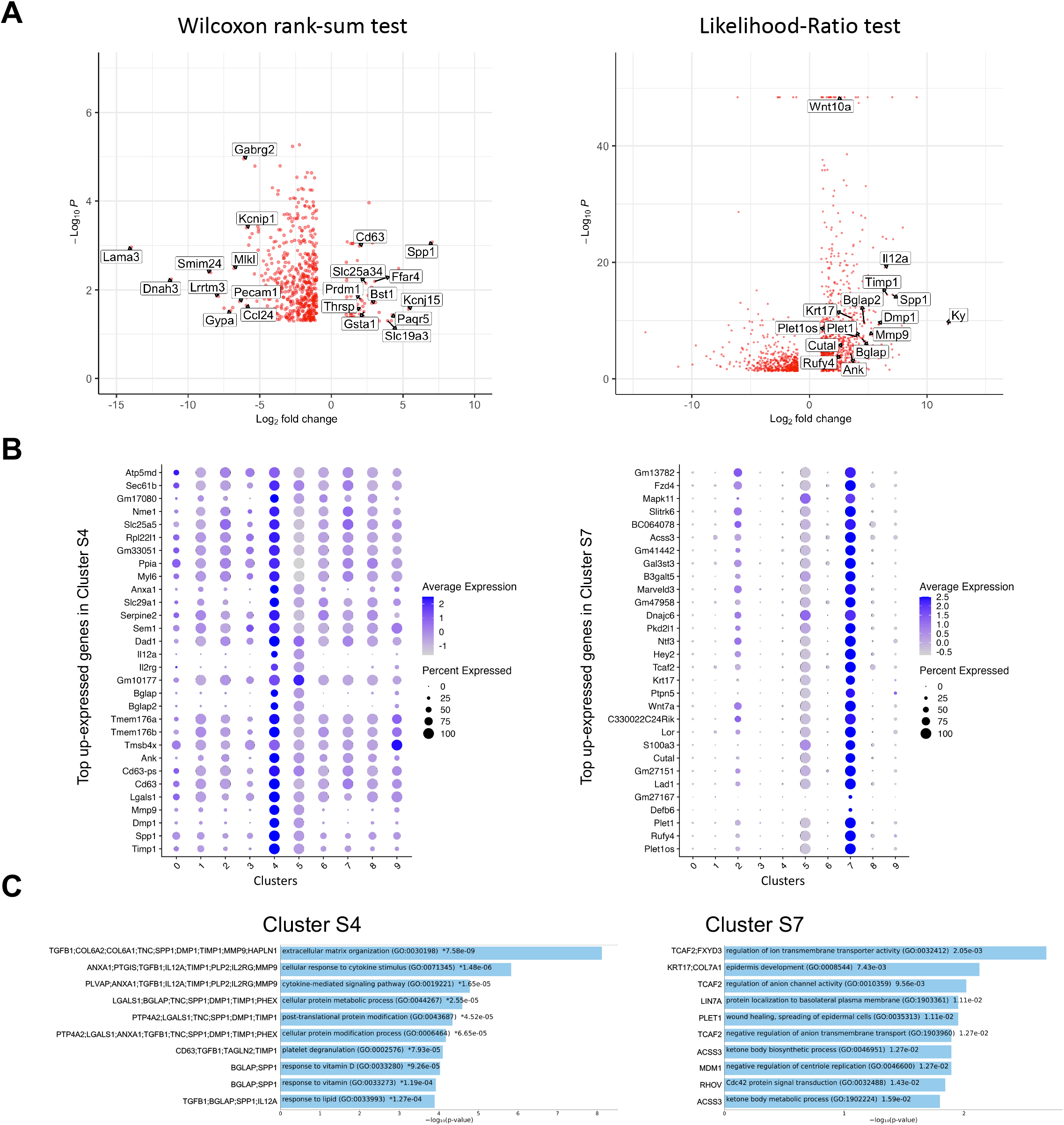
Analysis of DEGs in CA-ERBB2 cells and top up-expressed genes identified in cluster S4 and S7. (A) Volcano plots showing the distribution of gene expression fold changes and *p* values for DEGs identified in Wilcoxon rank-sum test (left) and Likelihood-Ratio test (right). Top up- and down-regulated genes with mapped IDs in the PANTHER classification system are indicated in the Wilcoxon rank-sum test volcano plot. Selected genes of interest are indicated in the Likelihood-Ratio test volcano plot. Complete list of genes is provided in S7 Data and S8 Data. (B) Dot plots showing the expression distribution of the top 30 genes of cluster S4 and S7 among all clusters. Complete list of up-expressed genes is provided in S9 Data. (C) Gene Ontology (GO) term enrichment analysis for up-expressed genes of cluster S4 and S7. Bar charts shows the top 10 enriched GO terms of biological process for S4 and S7, along with corresponding *p*-values (< 0.05). An asterisk (*) next to a *p*-value indicates the term also has a significant *q*-value (<0.05). The *y*-axis represents biological process, and the horizontal axis represents the number of genes, which are listed on the left side of the graph. Complete list of terms is provided in S12 Data.

In both analyses, *Spp1* was among the most up-expressed genes (132-fold change) in about 80% of CA-ERBB2 cells, in comparison to expression of *Spp1* in 55.4% of Control cells. *Spp1* will be more fully described later. *Lama3* was among the most down-regulated genes in CA-ERBB2 cells in both tests. It was detected in 3.4% of CA-ERBB2 cells, compared to 23.2% of Control cells. LAMA3 is a subunit of laminins, which mediate the attachment, migration and organization of cells by interacting with integrin receptors and other extracellular matrix (ECM) components (26, 27). In the LR test, *Kyphoscoliosis peptidase* (*Ky*) gene is top differentially expressed (3530-fold change), but only in about 7.3% of CA-ERBB2 cells, in comparison to 21.5% of Control cells. KY’s proposed function is maintenance of cytoskeleton and neuromuscular junction (28). These data suggest that some CA-ERBB2 cells modify the local environment, potentially affecting the behavior of neighboring cells.

To better understand transcriptional heterogeneity among CA-ERBB2 cells in clusters S4 and S7, we identified up-expressed genes in each cluster described in Fig 1C, and then examined activated pathways against the Gene Ontology (GO) biological process database. For each of 10 clusters, we analyzed gene expression in comparison to remaining clusters. Using the criteria of an adjusted *p*-value of <0.05 and a difference in levels of expression greater than 2-fold, we found 90 genes up-expressed in cluster S4, and 1102 genes in cluster S7. The expression of thirty top up-expressed genes identified in clusters S4 and S7 in comparison to other clusters is shown in Fig 3B. For the remaining clusters, which are composed of Control and CA-ERBB2 cells (except cluster S5), we found that S0 had 1125 up-expressed genes, S1 – 810, S2 – 731, S3 – 359, S5 – 3722, S6 – 484, S8 – 426, and S9 – 144 (S9 Data). The top 10 up-expressed genes in each cluster were then used to generate heatmaps (S10 Fig).

The analysis revealed that among top up-expressed genes in cluster S4 are DEGs identified in Wilcoxon rank-sum test (*Spp1, CD63, Tmsb4x;* Fig 3A; S7 Data), and in the LR test (*Spp1, Timp1, Dmp1, Mmp9, CD63;* Fig 3A; S8 Data). According to the heatmap, S4 was not related to any other cluster (S10 Fig). In contrast, cluster S7, comprised of CA-ERBB2 cells only, had top up-regulated genes in common with cluster S2, which was mostly Control cells. This suggests that CA-ERBB2 cells of cluster S7 originated from Control cells from cluster S2. Among the top up-expressed genes of S7 are DEGs identified in the LR test (*Plet1os, Rufy4, Plet1, Cutal*) (Fig 3A; S8 Data).

With respect to clusters comprised of both CA-ERBB2 and Control cells, some transcriptional relationships were also evident (S10 Fig). For example, cluster S6 had similarities to cluster S1, which is in proximity on the UMAP analysis. There were no marker genes up-expressed in cluster S0, S3 and S5. Surprisingly, in cluster S5 composed mostly of Control cells some level of up-regulation was observed for most of the up-expressed genes identified in other clusters, with exception of genes in cluster S8. Finally, clusters S8 and S9 are represented by small number of cells with up-expressed genes generally not found in other clusters, indicating that they are transcriptionally distinct populations. This analysis underscores the diversity of responses by SCs to CA-ERBB2 expression.

### CA-ERBB2 cells express genes involved in modulation of extracellular matrix

Gene set enrichment analysis against GO biological process was performed for up-expressed genes of cluster S4 and S7 to identify molecular pathways responding to ERBB2 activation (Fig 3C). For other clusters, results with listed 10 top GO terms with significant *q*-value <0.05 are in S11 Fig, and complete list of terms for all clusters is provided in S12 Data. Among the most enriched annotations, both by adjusted *p*-value and number of genes involved, were annotations of genes linked to modulation of ECM. The distinctive cluster S4, formed entirely by CA-ERBB2 cells, showed various degrees of activation of markers involved in extracellular matrix organization (GO:0030198), and extracellular matrix disassembly (GO:0022617). Cluster S4 had also enriched genes associated with cellular response to cytokine stimulus (GO:0071345), cytokine-mediated signaling pathway (GO:0019221) and regulation of integrin-mediated signaling pathway (GO:2001044). We note that clusters S1 and S6 were also distinguished by up-expressed genes involved in extracellular matrix organization and disassembly.

With respect to the distinctive cluster S7 of CA-ERBB2 cells, GO biological process analysis did not reveal terms with *q*-value <0.05 (Fig 3C). However, among terms with significant *p*-value (<0.05), are terms associated with genes *Krt17* and *Plet1* listed as DEGs in the LR test (Fig 3a; S8 Data). Epidermis development (GO:0008544) is associated with *Krt17* and *Col7a1* genes. This term is in a parent-child relationship with the term related to HC differentiation (inner ear auditory receptor cell differentiation, GO:0042491). Moreover, *Plet1* is associated with wound healing, the spreading of epidermal cells (both GO:0035313 and GO:0044319), and negative regulation of cell-matrix adhesion (GO:0001953) (S12 Data).

In clusters S1, S2 and S6, CA-ERBB2 cells were intermingled with Control cells suggesting that they have similar transcriptome. The identified GO terms for these clusters indicate that cochlear cells of P3 mice still undergo changes related to development of the organ of Corti (S11 Fig). For example, cells in cluster S1 were enriched of genes associated with sensory organ morphogenesis (GO:0090596), collagen fibril organization (GO:0030199), regulation of endothelial cell migration (GO:0010594), endodermal cell differentiation (GO:0035987), and skeletal system development (GO:0001501). In cluster S2, cells showed up-regulation of genes involved in epithelial cell development (GO:0002064), regulation of transcription involved in cell fate commitment (GO:0060850), and NOTCH signaling pathway (GO:0007219). In cluster S6, enriched genes are associated with sensory perception of mechanical stimulus (GO:0050954), and sensory perception of sound (GO:0007605).

Cells in cluster S8 were enriched of terms associated with cellular division. Analysis of transcriptomes for cell cycle-specific gene expression confirmed that cells in cluster S8 are in S and G2/M phase (S11 Fig and S13 Fig). The analysis also revealed that whereas most cells across the conditions were in G1 phase, clusters S4 and S0 also showed enrichment of cells in phase S and G2/M, indicating some cells prepared for, or underwent cell division (S13 Fig). In cluster S8, both CA-ERBB2 cells and Control cells showed markers for phase S and G2/M, however in cluster S0 this population was mostly represented by CA-ERBB2 cells.

### Top up-regulated genes in cluster S4 are in close network of predicated protein-protein interactions

As one goal of this study was to identify candidate secondary signaling pathways that could act downstream of ERBB2 signaling, we focused the next analysis on the new population of CA-ERBB2 cells that formed a transcriptionally distinct cluster S4. Top up-regulated genes *Spp1, Timp1, Mmp9* and *Dmp1*, are found in GO annotations associated with modulation of ECM and cytokine response (Fig 3C). Using STRING software, we analyzed protein-protein interaction network for SPP1, TIMP1, MMP9 and DMP1. The results are visualized in Fig 4A, and scores for predicted interactions are summarized in Table 1. The analysis shows that interactions between SPP1, TIMP1, MMP9 and DMP1 have association scores in the high and the highest confidence range (scores above 0.7 and 0.9, respectively). Increasing the network for additional five proteins revealed CD44 as protein with close association (Fig 4B), with the highest confidence score (above 0.9) calculated for SPP1, TIMP1, and MMP9, indicating likely true associations. Additional five proteins added to the network revealed two proteins that were found as differentially expressed between CA-ERBB2 and Control cells (Fig 4C). These proteins were CD63 and HPX, and both have association scores in the highest confidence range (above 0.9) with TIMP1; whereas HPX has also association score in the high confidence range (above 0.7) with MMP9.

**Fig 4.**
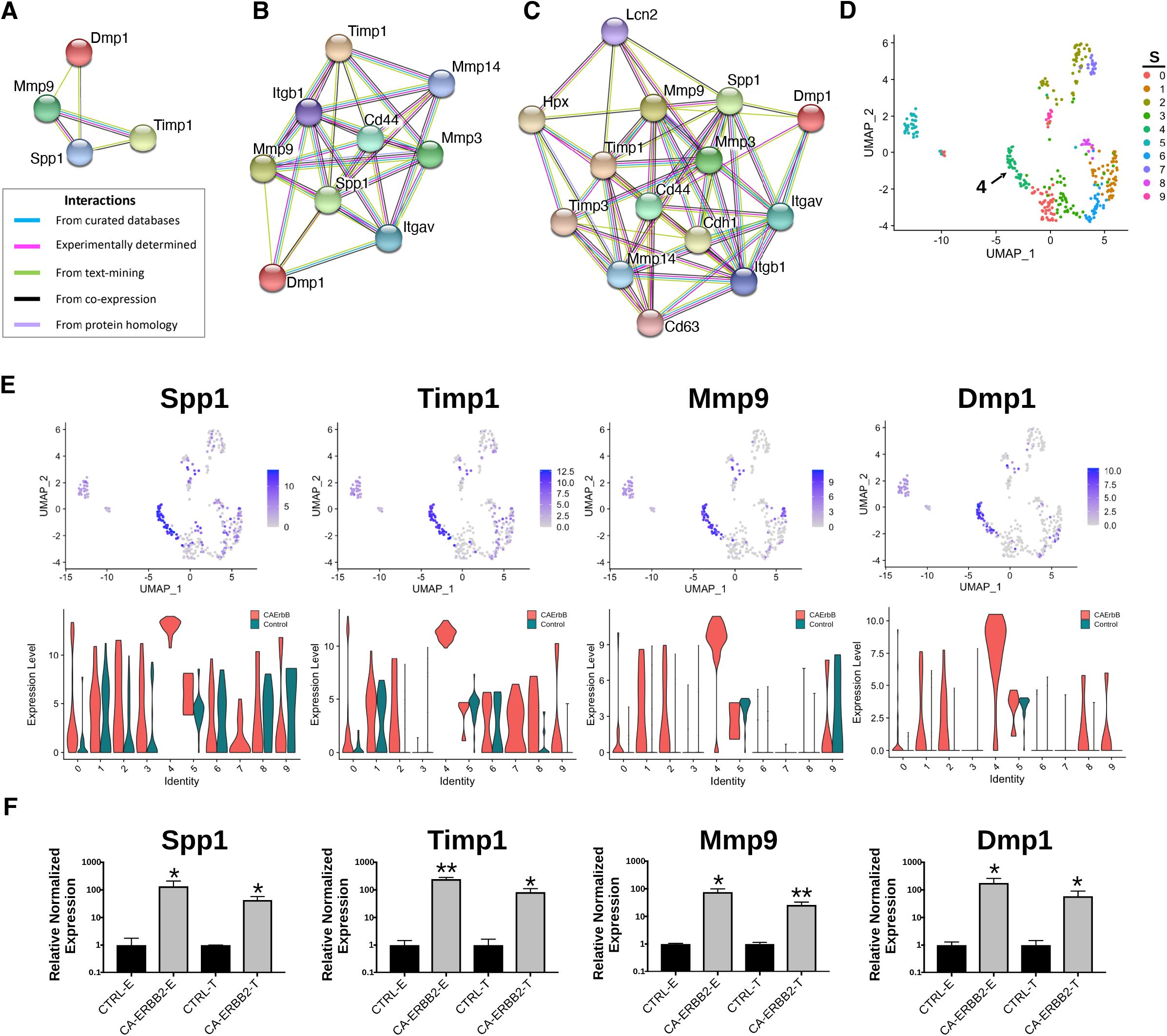
STRING protein-protein interaction network of four top up-regulated genes in cluster S4. (A) Protein-protein interaction network connectivity among SPP1, TIMP1, MMP9 and DMP1. In (B) and (C) protein-protein networks are shown with increasing number of closely associated genes. An edge was drawn with colored lines representing a type of evidence used in predicting the associations. A purple line indicates experimental evidence; a light green line indicates text-mining evidence; a light blue line indicates database evidence; a black line indicates co-expression evidence; and a light violet line indicates protein homology. The score interactions are summarized in Table 1. In (D), UMAP plot with marked cluster S4 enriched in cells expressing *Spp1, Timp1, Mmp9* and *Dmp1* in response to induction of ERBB2 signaling. In (E), UMAP plots and split violin plots showing up-regulated expression of *Spp1, Timp1, Mmp9* and *Dmp1* in CA-ERBB2 compared to Control cells in different clusters. (F) RT-qPCR validation of scRNA-seq analysis performed on GFP+ FACS-sorted cells from P3 cochleae. The level of gene expression in CA-ERBB2 sample is presented as ΔΔCt value (± SD) relative to Control (CTRL) normalized against the expression of the two reference genes *Eef1a1* (E) and *Tubb4a* (T) (n=3). Significance (\**p*<0.05; ** *p*<0.01) was determined by unpaired *t*-test (one-tailed) analysis.

**Table 1.**
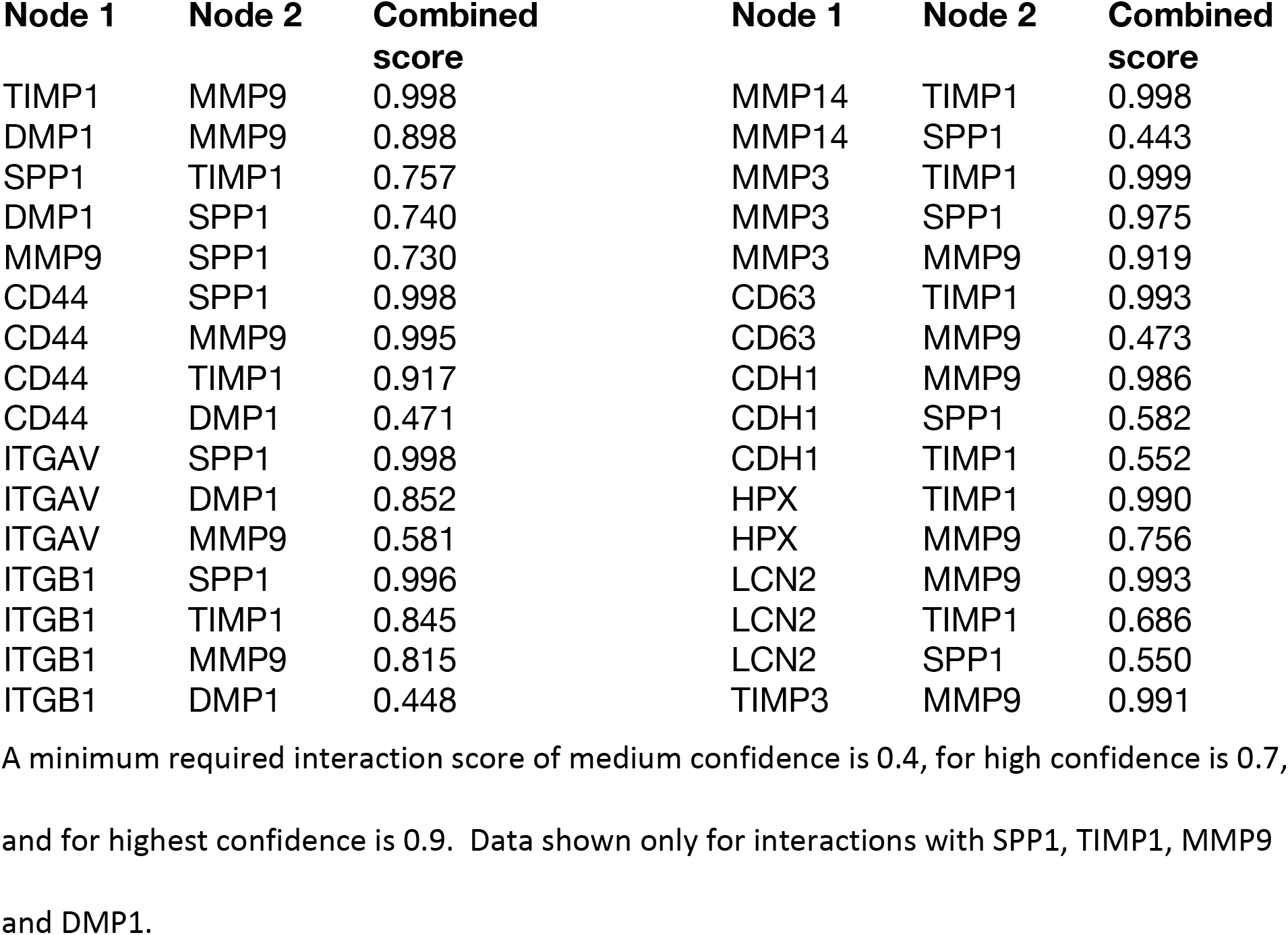
The combined score of interactions between pair of proteins in STRING database analysis.

| **Node 1** | **Node 2** | **Combined score** | **Node 1** | **Node 2** | **Combined score** |
| --- | --- | --- | --- | --- | --- |
| TIMP1 | MMP9 | 0.998 | MMP14 | TIMP1 | 0.998 |
| DMP1 | MMP9 | 0.898 | MMP14 | SPP1 | 0.443 |
| SPP1 | TIMP1 | 0.757 | MMP3 | TIMP1 | 0.999 |
| DMP1 | SPP1 | 0.740 | MMP3 | SPP1 | 0.975 |
| MMP9 | SPP1 | 0.730 | MMP3 | MMP9 | 0.919 |
| CD44 | SPP1 | 0.998 | CD63 | TIMP1 | 0.993 |
| CD44 | MMP9 | 0.995 | CD63 | MMP9 | 0.473 |
| CD44 | TIMP1 | 0.917 | CDH1 | MMP9 | 0.986 |
| CD44 | DMP1 | 0.471 | CDH1 | SPP1 | 0.582 |
| ITGAV | SPP1 | 0.998 | CDH1 | TIMP1 | 0.552 |
| ITGAV | DMP1 | 0.852 | HPX | TIMP1 | 0.990 |
| ITGAV | MMP9 | 0.581 | HPX | MMP9 | 0.756 |
| ITGB1 | SPP1 | 0.996 | LCN2 | MMP9 | 0.993 |
| ITGB1 | TIMP1 | 0.845 | LCN2 | TIMP1 | 0.686 |
| ITGB1 | MMP9 | 0.815 | LCN2 | SPP1 | 0.550 |
| ITGB1 | DMP1 | 0.448 | TIMP3 | MMP9 | 0.991 |

A minimum required interaction score of medium confidence is 0.4, for high confidence is 0.7, and for highest confidence is 0.9. Data shown only for interactions with SPP1, TIMP1, MMP9 and DMP1.

SPP1 and DMP1 are small integrin binding n-linked glycoproteins (SIBLINGs), which are secreted ligands for CD44 and integrin αvβ3 receptors. They promote survival, proliferation, differentiation, and migration in different systems (29–37). TIMP1 inhibits MMP9, a metallopeptidase that modulates both CD44 (38, 39) and NOTCH signaling (40). The CD44 receptor has a role in cell survival and proliferation in cancer development, but a recent finding also highlights its important role in tissue regeneration and wound healing (33, 41–43). Notably, CD44 is endogenously expressed on OPCs in mice in adulthood (44).

Analysis of UMAP and split violin plots for *Spp1, Timp1, Mmp9* and *Dmp1* indicates that up-expression of these genes is observed in CA-ERBB2 cells for most clusters, with significant enrichment in cluster S4 (Fig 4E). Control cells express lower levels of *Spp1* and *Timp1* genes, particularly in cluster S1, S5 and S6. For genes *Mmp9* and *Dmp1*, almost no expression is detected in Control cells, except for cluster S9 and S5. To validate the scRNA-seq analysis, we performed reverse transcriptase quantitative polymerase chain reaction (RT-qPCR) on FACS-sorted cochlear GFP+ cells collected from CA-ERBB2 and Control neonatal mice. Results of RT-qPCR analysis are shown in Fig 4F. They confirm that activation of ERBB2 signaling in CA-ERBB2 cells drove significant up-regulated expression of genes *Spp1, Timp1, Mmp9* and *Dmp1*, and also other genes identified as up-expressed in CA-ERBB2 cells of cluster S4 (*Bglap, Bglap2, Ptgis, Cd63, Il12a, Ank*) (S14 Fig and Fig 3B).

### Induction of ERBB2 signaling in deafened adult mice results in up-regulation of *Spp1 and Timp1*, activation of CD44 receptor, and formation of cellular clusters in the cochlear duct

To determine if the SIBLING cluster could be induced at adult stages, we used immunostaining on sections of cochlea from adult young mice after both damage-induced hearing loss and CA-ERBB2 induction. Male and female young adult mice with Control and CA-ERBB2 genotype were exposed to traumatic noise as described in the Methods (Fig 5A). At 3 days post noise exposure (3 DPN), the mice from both genotypes were randomly assigned to 2 groups, and treated with doxycycline (DOX) injection to initiate CA-ERBB2 expression; or with saline injection to use as reference. Mice were euthanized at 5 DPN, and their cochlea were dissected and processed for immunostaining with antibodies specific for SPP1, TIMP1, and the intracellular domain of the CD44 receptor.

**Fig 5.**
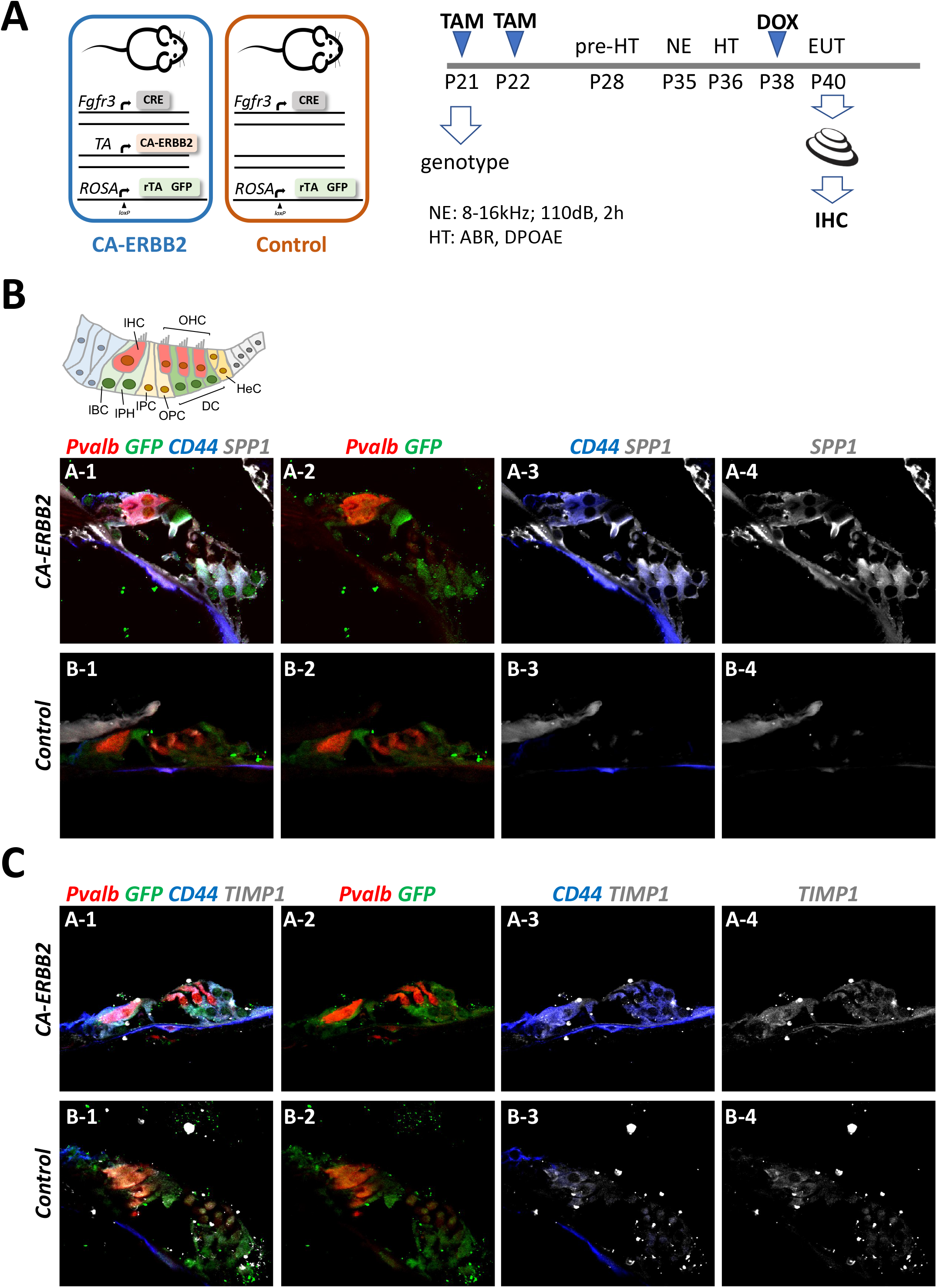
SPP1 and TIMP1 together with CD44 signaling are up-regulated in cochlear cells two days after CA-ERBB2 induction in noise exposed young adults. (A) Timeline of noise exposure (NE), GFP activation (TAM), CA-ERBB2 activation (DOX), and hearing tests (HT) in young adult mice. Mice were exposed to 110 dB octave band noise for 2 hours, and 3 days later injected with doxycycline (DOX) to induce CA-ERBB2 expression. HT were performed 1 DPN to confirm hearing loss. Immunostaining of cochlea sections revealed detection of SPP1 (B) and TIMP1 (C) together with activated CD44 receptor in CA-ERBB2 animals. Immunodetection for hair cells (PVALB, red), and cells competent to express CA-ERBB2 (GFP, green), are shown in panels 1 and 2. Immunodetection of activated CD44 receptor (blue) is shown in panels 1 and 3. Immunodetection of SPP1 and TIMP1 are shown in panel 1, 3 and 4 (gray).

As shown in Fig 5B and 5C, SPP1 and TIMP1 were up-expressed in cochlear sections from CA-ERBB2 animals treated with DOX. The fluorescent immunoreactivity for SPP1 corresponded to GFP+, including cells near OHC that correspond to DC, and cells near IHC corresponding to phalangeal cells and border cells. Similarly, TIMP1 was detected in GFP+ cells, although in sections from Control animals the immunoreactivity for TIMP1 was evident also in cells corresponding to inner phalangeal cells and border cells. Staining for the CD44 intracellular domain showed clearly up-regulation of CD44 signaling in GFP+ cells from CA-ERBB2 animals treated with DOX. The CD44 immunoreactivity was also observed in interphalangeal cells, DC and also in cells corresponding to HeC or Claudius cells. Interestingly, CD44 activation in cells corresponding to HeC or Claudius cells was also evident in sections from Control animals. This is consistent with previous report describing CD44 expression to increase in postnatal mice by P7 in the OPCs, Claudius cells and in a small number of GER cells (44).

Additional mice treated with traumatic noise, EdU, and doxycycline were euthanized at 7 DPN, and analyzed for GFP expression and additional markers (Fig 6). Here we found that aggregated GFP+ and GFP-negative cells were located in the endolymphatic duct of CA-ERBB2 cochlea. Phosphorylation of ERBB2 was seen in scattered cells, and did not precisely map onto GFP expression as observed at earlier time points(22). Mitotic figures labeled with EdU were evident in these aggregates (Fig 6B’), as were scattered cells labeled with activated CASP3 (Fig 6C’). Scattered cells within aggregates expressed sensory markers, including MYO7 (Fig 6B’) and the SC marker TAK1 (Fig 6D’). They were largely absent of SOX2 (Fig 6A’) and PVALB (Fig 6D’) expression. Some aggregates had smooth, compacted multicellular regions that stained brightly for phalloidin. Aggregates were observed in four out of five cochleae with CA-ERBB2 induction. In Control cochleae, only very small clusters of cells were observed floating in the cochlear duct, none of which expressed GFP or incorporated EdU (Fig 6E, E’, arrows).

**Fig 6.**
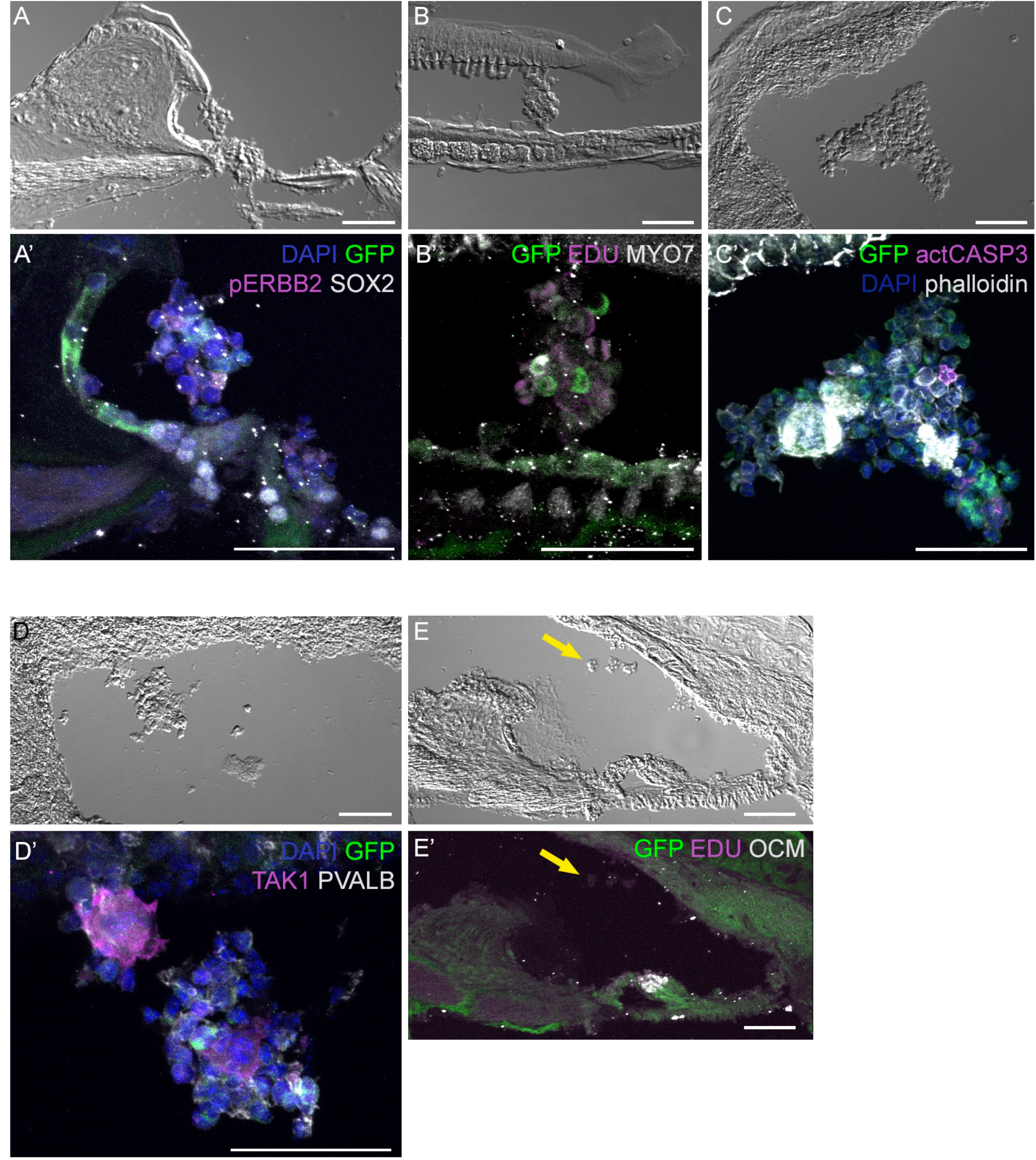
Four days after CA-ERBB2 induction, aggregates of sensory cells can be found within the cochlear duct. Brightfield low power images are used to show the position of the cellular aggregates, while immunostaining reveals specific markers. (A) and (A’), GFP (green) and pERBB2 (magenta) are present in an aggregate attached to the spiral limbus, which has little SOX2 (white). (B) and (B’), A similarly localized aggregate harbors GFP+ cells (green) and rare MYO7+ cells (white), with EdU+ mitotic figures (magenta) in cells lacking GFP. (C) and (C’), An aggregate near the stria vascularis contains GFP+ cells (green), rare apoptotic cells marked with anti-activated CASP3 (magenta), and compacted epithelial cells revealed by phalloidin staining (white). (D) and (D’), Another aggregate near the stria vascularis harbors cells with GFP (green) and neighboring cells positive for the supporting cell marker TAK1 (magenta), but no PVALB+ hair cells. (E) and (E’), A Control mouse lacking the CA-ERBB2 transgene harbors only small clusters of cells (yellow arrow) without GFP expression.

## DISCUSSION

Previously, we found that SCs with activated ERBB2 influenced gene expression in their neighbors *in vivo* and promoted the differentiation of hair-cell like cells, suggesting that ERBB2 may initiate a signaling cascade important for HC regeneration (22). Here, single cell transcriptome analysis revealed the formation of new population among a subset of cells with activated ERBB2 signaling, which had differentiated significantly from the parental Control cells. Possibly derived from OPCs, these cells also express LGER-specific genes. They comprise most of the cells expressing *Spp1*, identified in our studies as one of the most highly activated genes in response to ERBB2 signaling. Importantly, in this novel population we see expression of the other three top up-regulated genes, *Mmp9, Timp1* and *Dmp1*. Together with *Spp1*, they are associated with a signaling network involving CD44 and integrin αvβ3 receptors. Gene ontology enrichment analysis revealed that these genes modulate both the ECM and cytokine responses, including Notch signaling. We confirmed that SPP1, TIMP1 and CD44 protein are also up-regulated in adult cochlear SCs after CA-ERBB2 induction. Subsequently, we observed the formation of floating aggregates of cochlear sensory cells. The fate and potential of these cellular aggregates are not known.

SPP1 and DMP1 are secreted glycoproteins that act as ligands for CD44 and integrin αvβ3 receptors. They are members of the small integrin-binding ligand n-linked glycoprotein (SIBLING) family. SPP1 promotes survival, proliferation, differentiation, adhesion, and migration of numerous cell types (29–35, 37). While DMP1 function in injury and wound healing has not been yet determined, it is significantly up-regulated in a number of cancerous tissues (45, 46). SPP1 may function in the neuronal differentiation of auditory neurons during inner ear development (47). Moreover, it is detected in membranous labyrinth of the adult mammalian cochleae, particularly in the utricle, where it is currently used as a marker of type I HC (48–51).

The most significant aspect of up-regulation of SPP1, MMP9 and TIMP1 in response to ERBB2 signaling, is that they promote wound healing and regeneration after injury in other systems (33, 41–43). SPP1 is well known to mediate bone and muscle regeneration (52–56), and it is also involved in promoting proliferation and regeneration in nervous system (57–59). In recent studies, Wang et al. demonstrated that binding of SPP1 with CD44 and integrin αvβ3 is important in the Schwann cells function in regenerating nerves, by promoting proliferation and survival after peripheral nerve injury (60). Another study by Powell and colleagues showed that SPP1 together with MMP9 acts through CD44 receptor to mediate synaptogenesis after central nervous system (CNS) insult (61).

TIMP1 is an inhibitor of MMP9 and the balance between them is a key regulator of the signaling network in the injured nerve (43). The significance of MMP9/TIMP1 balance is not well understood, but it was also demonstrated to regulate healing process of burned fetal skin (62). An increased MMP9/TIMP1 ratio promotes the degradation of ECM to allow cellular migration, while a reduced MMP9/TIMP1 ratio drives reconstruction of the ECM during healing process (62). TIMP1 inhibition of MMP9 could explain the widespread downregulation of SOX2 after CA-ERBB2 signaling in vivo (22), as MMP9 is necessary for NOTCH signaling (63), and NOTCH is necessary and sufficient to promote SOX2 expression (64). TIMP1 may also act as cytokine independently of MMP9 inhibition to promote cell survival and proliferation (65). In hematopoietic stem and progenitor cells, TIMP1 promotes migration, adhesion and survival by binding to CD63-integrin β1 receptors complex (66). Our scRNA-seq analysis revealed CD63 as DEG and its up-regulation was mostly enriched in cluster S4 formed by CA-ERBB2 cells (S7 Data and S8 Data).

Up-regulation of SPP1, MMP9, TIMP1 and DMP1 in response to ERBB2 activation suggests that its downstream signaling involves CD44 receptor and potentially also integrin receptors. The relationship between ERBB2 and CD44 was previously described for maintaining neuron–Schwann cell interactions in early rat neonatal nerves development (67). In particular, CD44 significantly enhanced neuregulin-induced ERBB2 phosphorylation and ERBB2–ERBB3 heterodimerization. CD44 was also found to interact with ERBB2 receptor (67, 68). Importance of CD44 receptor’s network in cells survival, proliferation and regeneration described for Schwann cell and in other tissues, suggests that some of CA-ERBB2 effects involve signaling through CD44 receptor.

Lower flexibility of the epithelium associated with cell-cell junction was proposed as one of the mechanisms responsible for decreased regenerative capacity of the adult mammalian cochlea (69). Our results indicate that ERBB2 signaling promotes modulation of ECM that would allow increased flexibility in the organ of Corti. The combination of EGF and GSK3 inhibitors were recently reported to deplete E-cadherin in tight junctions of the adult mammalian cochlea (70). In addition to up-regulated genes associated with ECM disassembly in cluster S4, there is also up-regulated *Plet1* gene in CA-ERBB2 cells from cluster S7, which is associated with negative regulation of cell-matrix adhesion (GO:0001953). *Plet1* is listed in GO terms associated with spreading of cells and wound healing (GO:0035313, GO:0044319). Thus, we would predict that its up-regulation would allow for increased local movements of cells.

Importance of ERBB2-mediated ECM modulation for HCs regeneration is also supported by recent studies indicating that ECM and integrin receptors are induced in early neurosensory development and cell fate determination in the human fetal inner ear (71). We speculate that enforced activation of signaling pathways through ERBB family receptors might allow for changes in the environment of the organ of Corti that promote the proliferation of SCs and permit the regeneration of HCs. Previous studies have shown little or no stem cell activity in the adult mammalian cochlea (16, 72). This contrasts to robust stem-like capacity in isolated cells from the adult utricle (16, 73), which correlates with limited regeneration potential in that organ (74). These data may be interpreted to mean that cochlear stem cells, perhaps resident in GER (75), are lost during maturation. It is also possible that their requirements for survival or for identity maintenance differ. For example, they may require accessory cells to maintain their microenvironmental niche. Alternatively, the forced expression of CA-ERBB2 signaling might promote novel behaviors not otherwise seen in cochlear SCs.

In conclusion, we report that enforced signaling of ERBB2 in a minority of cochlear SCs, possibly OPCs, drives a novel differentiation response in the neonatal mouse cochlea. These cells induce expression of a cluster of genes involved in signaling, wound healing, and migration. We confirm that at least two of these genes, SPP1 and CD44, are also up-regulated in adult cochlear cells after CA-ERBB2 induction. CA-ERBB2 induction also correlates with the generation of cellular aggregates in the cochlear duct. These aggregates contain, but are not limited to, fate-mapped CA-ERBB2 cells. Their potential, effects, and fate remain to be determined.

## METHODS

### Ethical Approval

All experiments using animals were approved in advance by the University Committee on Animal Research (UCAR) at the University of Rochester, protocol number 2010-011, PI Patricia White, and by the Animal Care and Use Review Office (ACURO) of the Department of Defense.

### Mice

The following mouse strains were used. Fgfr3-iCre (23) was a kind gift from Dr. Jian Zuo. TetON-CA-ErbB2 (76) and CBA/CaJ, were purchased from Jackson Laboratories. ROSA-floxed-rtTA/GFP (77) was a kind gift from Dr. Lin Gan. TetON-CA-ErbB2 mice harbor a mutated *ErbB2* transgene encoding a constitutively active ERBB2 protein (CA-ERBB2), which does not require ligand binding or heterodimerization with other ERBB partners for active signaling. Fgfr3-iCre and TetON-CA-ERBB2 mice were crosses to generate double heterozygous transgenic mice that were subsequently used as breeders. In a second breeding, double heterozygous were crossed with homozygous ROSA-floxed-rtTA/GFP mice.

### Genotyping

For scRNA-seq experiments and data validation by RT-qPCR, generated pups were genotyped at P0 to identify triple-transgenic mice that harbor *Fgfr3-iCre*, ROSA-floxed-rtTA/GFP, and TetON-CAERBB2 (named here as CA-ERBB2). Oligonucleotides used in genotyping are provided in S15 Table. Littermates with both *Fgfr3-iCre* and ROSA-floxed-rtTA/GFP were used as Controls.

### Noise exposure

For scRNA-seq data validation in young adults exposed to noise, both male and female mice were used equally. Mice harboring the *Fgfr3-iCre* and *CA-Erbb2* transgenes have been backcrossed to CBA/CaJ for at least four generations, whereas the ROSA-rtTA-GFP line is congenic on C57BL6/J. Thus, the experimental mice are hybrid CBA/C57 mice, whose noise damage characteristics are similar to a pure CBA/CaJ response (78). Three-week-old mice were genotyped and weaned. Tamoxifen injections (75 mg/kg) were performed at P21, P22 and P23. At P28 their hearing was tested by auditory brainstem response (ABR) and distortion product acoustic emissions (DPOAE) (79). The mice were then divided into DOX exposed and control conditions. Mice experienced a noise exposure of 110 dB of 8-16 kHz octave band noise for 2 hours, as described previously (79). The next day (1 DPN), mice received hearing tests to confirm severe noise-induced hearing loss evident by threshold shift for at least 25 dB across five frequencies (8, 12, 16, 24 and 32 kHz). Two days later (3 DPN), the DOX group were injected with DOX (100 mg/kg), followed 30 minutes later by an injection of furosemide (400 mg/kg) (80). The control group received a saline injection, followed by furosemide. Two days after these injections (5 DPN), the mice were euthanized and their cochleae fixed overnight in fresh 4% paraformaldehyde. A second cohort of mice were treated identically, except that they also received 2 injections of 10 mg/kg 5-ethynyl-2’-deoxyuridine (EdU) from the Click-iT EdU Alexa Fluor 647 imaging kit (Invitrogen, Carlsbad, CA), the day after DOX injection (4 DPN) spaced 8 hours apart. These mice were euthanized on 7 DPN and their cochleae were fixed overnight in fresh 4% paraformaldehyde.

### Cell dissociation and FACS sorting

The iCRE activity was induced by tamoxifen injections (75 mg/kg) at P0/P1 to label supporting cells with GFP and rtTA. Doxycycline injections (100 mg/kg) were performed at P2 to drive CA-ERBB2 expression in GFP+ supporting cells. Cochlea from P3 pups were collected and incubated with 1 mL Accutase® solution (Innovative Cell Technologies) for 15 minutes at 37°C. To ensure a reproducible digestion, 2-4 cochlea were treated per tube. The Accutase solution was removed and cochlea were gentle washes in ice cold Ca2+, Mg2+ - free HBSS buffer. Organs in each tube were dissociated by trituration for 2-3 minutes. Cells were then filtered with a 40 μm cell strainer (BD Biosciences) and kept on ice until sorted. For the exclusion of dead cells and debris from the samples during sorting, a single-cell suspension was stained with DAPI. Sorting was performed at 4°C in a FACSAria II cell sorter (BD Biosciences) using a 130 μm nozzle.

Scatter discrimination was used to eliminate doublets and samples were negatively selected with DAPI to remove dead cells and debris. For scRNA-seq, single GFP+ cells were captured into 96-well plate with 11.5 μl lysis buffer and RNase Inhibitor without the CDS IIA Oligo (Takara Bio, Mountain View, CA), and stored at −80°C until used for construction of cDNA libraries. On average, 6 pups per genotype were processed for sorting single GFP+ cells into one 96-well plate. For data validation by RT-qPCR, GFP+ cells were captured into single tube with 100 μl lysis buffer containing Proteinase K and DNase (Bio-Rad). The sorted cells were then incubated for 10 min. in room temperature, followed by 5 min. at 37°C, and 5 min. at 75°C. The samples were stored at −80°C until RT-qPCR analysis.

### Construction of single-cell RNA-seq libraries

Single-cell RNA-seq libraries were constructed according to the SMART-Seq Single PLUS protocol (Takara Bio, Mountain View, CA). Frozen plates were thawed on ice. Next, cDNA was generated with the addition of the 3’ SMART-Seq CDS Primer II A and SMARTScribe II RT reagents. cDNA was amplified via 20 cycles of PCR, purified, and assessed for quantity and quality by Qubit dsDNA assay (ThermoFisher, Waltham, MA) and Fragment Analyzer (Agilent, Santa Clara, CA) analysis, respectively. For each well, 1 ng amplified cDNA was fragmented and prepared for library construction with the addition of stem-loop adapters. Final library amplification and indexing was performed through 16 cycles of PCR amplification. Following bead purification, individual library profiles were assessed by Fragment Analyzer and quantified by Qubit dsDNA assay. Libraries were diluted, pooled in equimolar amounts, and sequenced on a NovaSeq S1 flowcell (Illumina, San Diego, CA) to generate an average of 10 million reads per cell.

### Data preprocessing, clustering, and visualization

Raw reads generated from the Illumina basecalls were demultiplexed using bcl2fastq version 2.19.1. Quality filtering and adapter removal are performed using FastP version 0.20.1 with the following parameters: “--length_required 35 --cut_front_window_size 1 -- cut_front_mean_quality 13 --cut_front --cut_tail_window_size 1 --cut_tail_mean_quality 13 -- cut_tail -y -r”. Processed/cleaned reads were then mapped to the Mus musculus reference genome (GRCm38 + Gencode-M25 Annotation) using STAR_2.7.6a with the following parameters: “—twopass Mode Basic --runMode alignReads --outSAMtype BAM SortedByCoordinate – outSAMstrandField intronMotif --outFilterIntronMotifs RemoveNoncanonical –outReads UnmappedFastx”. Gene level read quantification was derived using the subread-2.0.1 package (featureCounts) with a GTF annotation file (Gencode M25) and the following parameters: “-s 2 -t exon -g gene_name”. The data generated by the sequencing was used to create a Seurat object using the Seurat package in R (v4.0.5 and v4.1.2). Quality control metrics were performed using standard Seurat (v4.0.4 and v4.1.0) protocols and included the identification of unique genes (“features”) per cell, the total number of molecules detected (“reads”), and the percent of mitochondrial genome contamination. The data was normalized. Clusters were identified with the Louvain method, using PCA and UMAP to reduce dimensionality and plot the cells. Ten clusters were identified using a dimensionality reduction of 1:30 within the Nearest-neighbor graph construction and UMAP generation, as well as a resolution of 2 within clustering. In addition to performing sample filtering and clustering, Seurat handled differential expression to determine genes up-regulated within clusters and across condition groups. enrichR (v3.0) was used to perform gene set enrichment on DEGs(81). A final report was rendered using Rstudio and rmarkdown (v2.10). Identity of genes was analyzed in the Gene Ontology resource (http://geneontology.org/) (81). GO biological process analysis was performed for up-expressed genes identified in each cluster. Since there were to many up-expressed genes for some of the clusters we limited the number of markers coming from specific clusters based off of log fold change. The log fold change threshold for cluster S7 was 4, for cluster S0 was 4, and for cluster S5 was 6. The log fold change threshold for the remaining clusters remained at 2. The protein interaction network modules were visualized using STRING network online analysis (http://string-db.org) (82).

### RT-qPCR analysis

Cell lysates prepared after sorting GFP+ cells into single tube were used in RT-qPCR analysis using SingleShot Two-Step RT-qPCR reagents (Bio-Rad) according to manufacture instruction. Each qPCR reaction was performed on cells lysates corresponding to 10 cells. Three technical replicates were used for each analyzed gene. Wells with primers but without cDNA sample were utilized for negative controls, while cell lysates not treated with reverse transcriptase were used as a control of genomic DNA contamination. qPCR reactions were performed using Bio-Rad CFX Thermal Cycler and the threshold of cycles (Ct) values was calculated with Bio-Rad CFX Manager Software. ΔΔCt method was used for calculating relative gene expression. *Eef1a1* and *Tubb4a* were used as reference genes to calculate differences in gene expression between Control and CA-ERBB2 cells (83). The Ct values of technical replicates were averaged and expression of the gene in CA-ERBB2 sample and Control sample was normalized to reference gene (“REF”) expression level within the same sample to determine ΔCt (Ct _(gene)_ – Ct _(REF)_). For each gene, analysis of relative gene expression was performed from at least three biological samples (three independent FACS sortings). The ΔCt for each biological replicate was exponentially transformed to the ΔCt expression (2^-ΔCt^) before averaging and determining the standard deviation (SD). The mean was then normalized to the expression of gene in Control sample to find ΔΔCt expression in CA-ERBB2 sample. Oligonucleotides used in RT-qPCR are provided in S15 Table.

### Immunochemistry

Fixed cochleae were decalcified in 100 mM EDTA in a saline solution at 4°C for 3 days. For cryosectioning, tissues were immersed in 30% sucrose in PBS overnight, embedded in OCT, frozen in liquid nitrogen, and cryosectioned at 20 microns. Sections were mounted on slide glass and kept at −30°C until use. Immunostaining of cryostat cochlear sections was performed to examine the expression of selected genes. The sections were permeabilized and blocked in PBS with 0.5% Triton X-100 and 5% donkey serum (TDB buffer) for 1 hour at 4°C, followed by incubation with primary antibodies in TDB at 4°C overnight. The sections were rinsed briefly 4 times in PBS, then incubated with secondary antibodies in TDB at 4°C overnight. The sections were rinsed briefly 4 times in PBS. The following primary antibodies were used: chicken anti-GFP (1:100; Abcam; AB_300798), mouse anti-PVALB (1:1000, EMD Millipore; MAB1572), goat anti-OPN (SPP1; 1:100; R&D Systems; AF808), goat anti-TIMP1 (1:100; R&D Systems; AF980), rabbit anti-CD44 (1:100; Abcam; AB157107), rabbit anti-pERBB2 (1:200; Santa Cruz; sc-12352), goat anti-SOX2 (1:500; Santa Cruz; sc-17320), rabbit anti-MYO7 (1:1000; Proteus; 25-6790), rabbit anti-activated CASP3 (1:500; R&D Systems, AF835), and rabbit anti-TAK1 (1:500; Thermo Fisher; 28H25L68). Secondary antibodies were purchased from Jackson Immunoresearch. EdU was revealed using the Click-iT EdU Alexa Fluor 647 kit (Invitrogen; C10340) following the manufacturer’s instructions. Imaging was performed using Confocal Microscope Olympus FV1000.

## DATA AVAILABILITY

All data supporting the findings of this study are available within the article and its Supplementary Information files. Single-cell gene expression data have been deposited in the Gene Expression Omnibus data repository under accession code: GSE202850. Original image files and qPCR data files are deposited at the Open Science Foundation, under the title of the paper.

## ACKNOWLEDGMENTS

We gratefully acknowledge Dr. Anne Luebke, who maintains the URMC Small Animal Auditory Testing Core; Dr. Jian Zuo for the Fgfr3-iCre mouse strain; Dr. Lin Gan for the ROSA-floxed rtTA/GFP mouse strain, and Dr. Amy Kiernan for advice on experimental interpretation.

## SUPPLEMENTARY FILES

**S1 Fig. Study design workflow for scRNA-seq analysis of transcriptomes of cochlear supporting cells expressing ERBB2.** (A) Breeding strategy to generate mice with CA-ERBB2 transgene inducible in cochlear supporting cells. (B) Timeline of tamoxifen (TAM) and doxycycline (DOX) injections in neonatal mice, and major steps in processing samples for scRNA-seq analysis. (C) Computational data set analysis of Control and CA-ERBB2 cells.

**S2 Fig. GFP labeling of supporting cells in the organ of Corti for FACS analysis.** The iCre gene under control of Fgfr3 promoter is mostly expressed in supporting cells, such as Deiter cells, Pillar cells and Hensen cells. Two injections of tamoxifen at P0 and P1 allow activation of iCRE and activation of GFP expression mostly in apical region of the organ of Corti. (A) Shown are confocal images of a whole mount preparations of the organ of Corti from apical, mid and basal portions. Samples were collected at P3. Both, IHC and OHC are labelled with antibodies to MYO-VIIA (MYO7). (B) Shown are flow cytometry plots with gating of cochlear cells dissociated from P3 pups for FACS purification of GFP+ cells. P1, initial gating of cells to exclude debris. P2 and P3, gating based on forward scatter height (FSC-H), forward scatter width (FSC-W) and side scatter width (SSC-W) for doublets discrimination to select single cells. P4, gating of live cells based on negative staining with DAPI. GFP+ population is selected for sorting from the DAPI-negative cells (P4).

**S3 Fig. Additional quality control metrics of scRNA-seq dataset.** (A) Violin plots showing similar distributions observed for cells from Control and CA-ERBB2 samples in percentage of reads from mitochondrial transcripts per cell. Most of the cells are identified as intact cells based on a low percentage of reads from mitochondrial transcripts, as anticipated by their exclusion of a cell impermeant dye. (B) UMAP plots showing distribution of percentage of mitochondrial gene (mt) expression in cells. (C) UMAP plot of CA-ERBB2 and Control cells in clusters for comparison.

**S4 Table. Cell-type specific markers.** Gene markers were selected based on (Kolla et al, 2020).

**S5 Fig. UMAP plots and violin plots for marker gene transcripts identifying cochlear cells among ten clusters**. Expression of gene markers was analyzed for Deiter cells, rows 1 and 2 (A), row 3 (B), Hensen cells (C), Inner Pillar cells (D), Outer Pillar cells (E), Inner Hair cells (F), Outer Hair cells (G), Lateral Greater Epithelial Ridge cells, group 1 (H), group 2 (I), group 3 (J), Medial Greater Epithelial Ridge cells (K), Inner Sulcus cells (L), Outer Sulcus cells (M), Interdental cells (N), and cells expressing Oc90 (O). Statistical analyses were performed with R version 4.0.5.

**S6 Fig. Graphical representation of statistically significant changes in gene expression between CA-ERBB2 SCs and Control SCs.** Graphs showing number of up- and down-regulated genes in CA-ERBB2 cells identified in Wilcoxon rank-sum test (**A**) and in Likelihood-Ratio test (**B**). In each group, protein coding transcripts are represented in grey, and non-coding RNA (ncRNA)/pseudogenes are represented in black. Numbers in bar indicate numbers of transcripts with mapped IDs (protein coding) and unmapped IDs (ncRNA/pseudogenes). On right, pie graphs for Wilcoxon rank-sum test (**A**) and Likelihood-Ratio test (**B**) showing the percentage of differentially expressed genes (DEG) found in up to 10% of cells, 10-20% of cells, 20-50% of cells, and in above 50% of cells.

**S7 Data. Differentially Expressed genes between CA-ERBB2 and Control cells in all data sets identified in the Wilcoxon rank-sum test**.

**S8 Data. Differentially Expressed genes between CA-ERBB2 and Control cells in all data sets identified in the Likelihood-Ratio test.**

**S9 Data. Up-Expressed genes by cell cluster.**

**S10 Fig. Heatmap depicting the top 10 genes expressed by 10 different clusters of cochlear SCs.** Clusters S4 and S7 are composed of CA-ERBB2 cells only. Cells from each cluster are arrayed along the horizontal axis and genes are arrayed along vertical axis. Clusters are distinguished by the color bars.

**S11 Fig. Top 10 significantly enriched Gene Ontology (GO) terms in different clusters.** For each cluster, bar chart shows the top 10 enriched GO terms of biological process along with corresponding p-values (< 0.05). An asterisk (*) next to a p-value indicates the term also has a significant q-value (<0.05). The y-axis represents biological process, and the horizontal axis represents the number of genes, which are listed on the left side of the graph. Complete list of terms for each cluster is provided in S12 Data.

**S12 Data. Gene set enrichment analysis.**

**S13 Fig. Cell cycle analysis.** Top left, UMAP plot showing distribution of cells with markers identifying cell cycle phases G1, G2/M and S. Top right, bar chart showing proportion of Control cells and CA-ERBB2 cells in cell cycle phase G2/M and S. Bottom bar charts show distribution of Control cells and CA-ERBB2 cells in G2/M and S phase by cluster.

**S14 Fig. Additional validation of up-regulated genes**. RT-qPCR was performed on GFP+ FACS-sorted cochlear cells from P3 pups. The level of gene expression in CA-ERBB2 sample is presented as ΔΔCt value (± SD) relative to Control (CTRL) normalized against the expression of the two reference genes, Eef1a1 (E) and Tubb4a (T) (n=3). Significance (*p<0.05; ** p<0.01; *** p<0.001) was determined by unpaired t-test (one-tailed) analysis. Below, UMAP plots and split violin plots showing up-regulated expression of Bglap, Bglap2, Ptgis, Il12a, CD63 and Ank in cells with activated CA-ERBB2.

**S15 Table. Oligonucleotides used in genotyping and RT-qPCR analysis.**

