## Supplementary figures and images for "Single cell RNA sequencing analysis of mouse cochlear supporting cell transcriptomes with activated ERBB2 receptor, a candidate mediator of cochlear regeneration mechanisms"

### S1 Fig

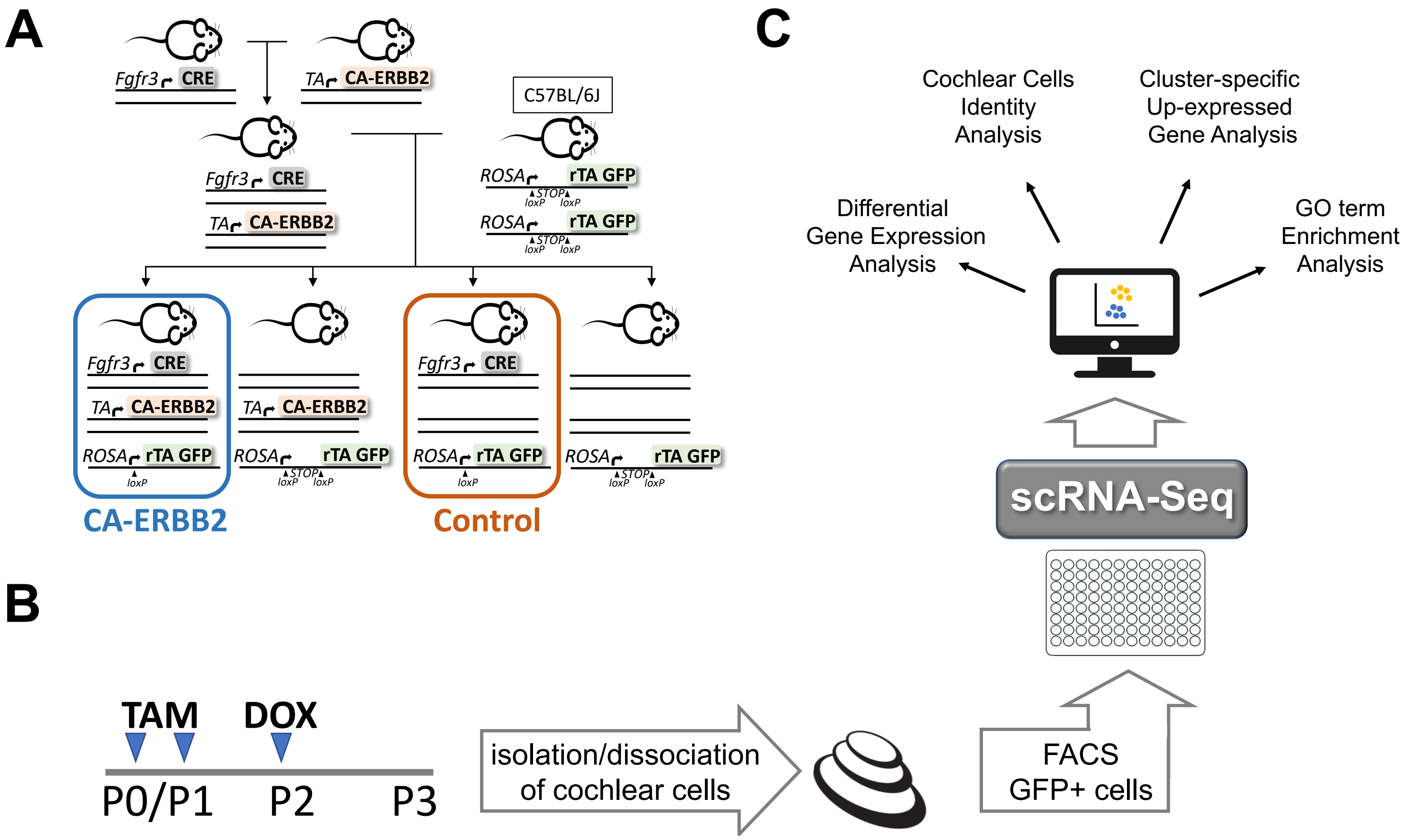

### S2 Fig

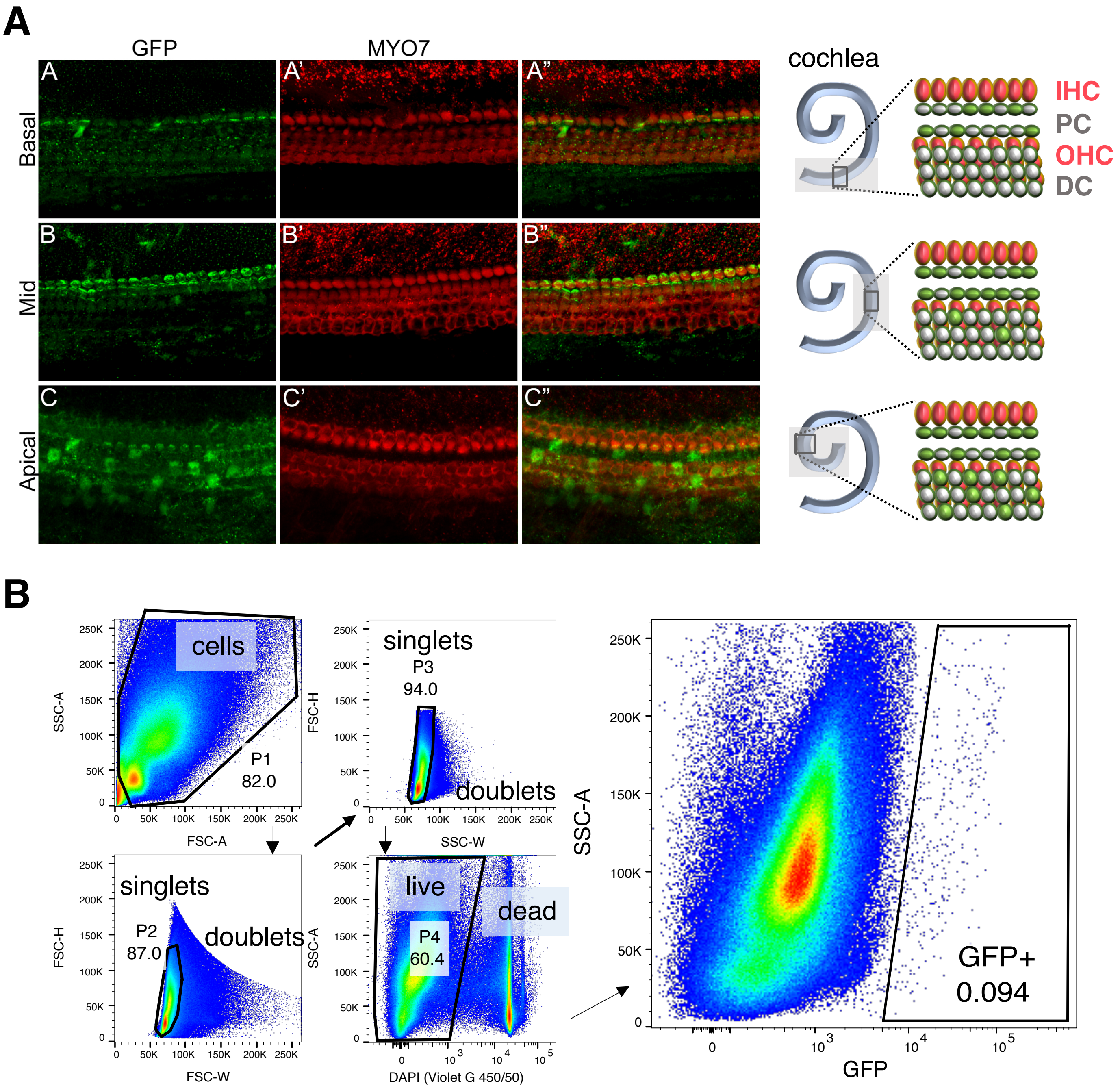

### S3 Fig

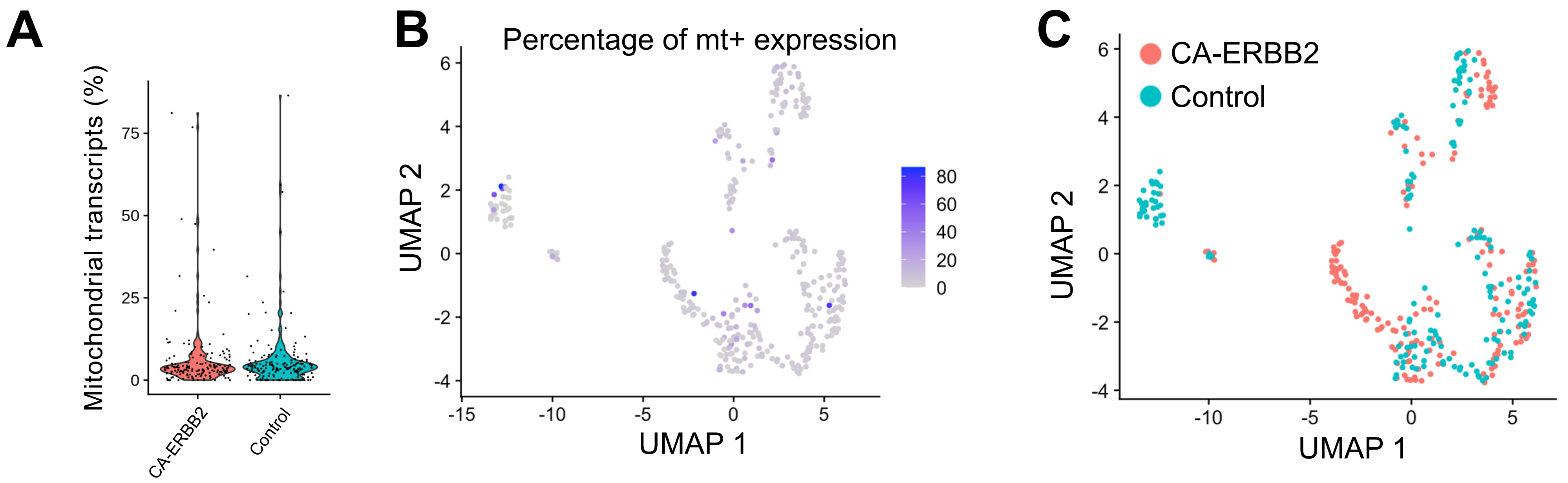

### S6 Fig

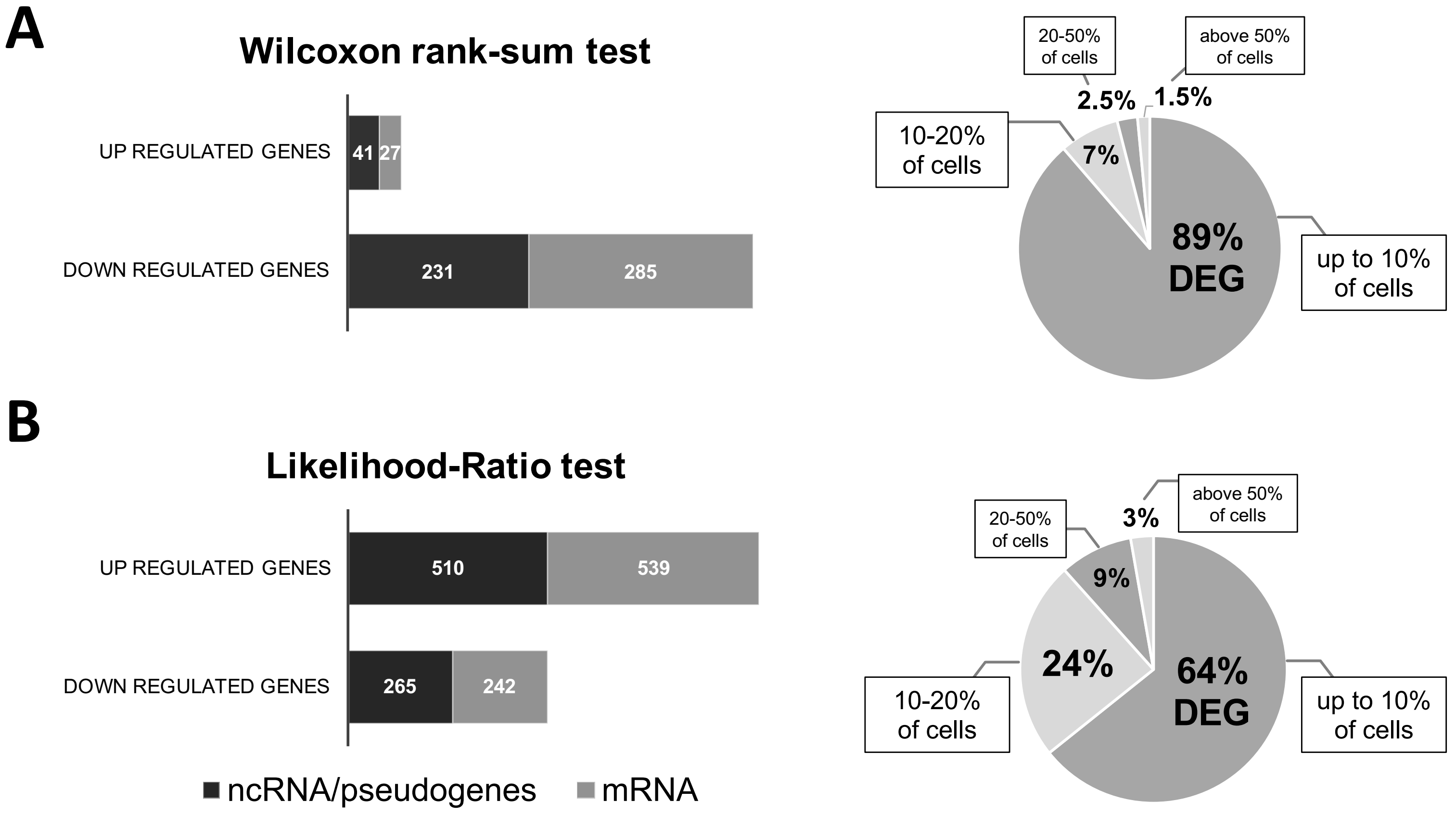

### S10 Fig

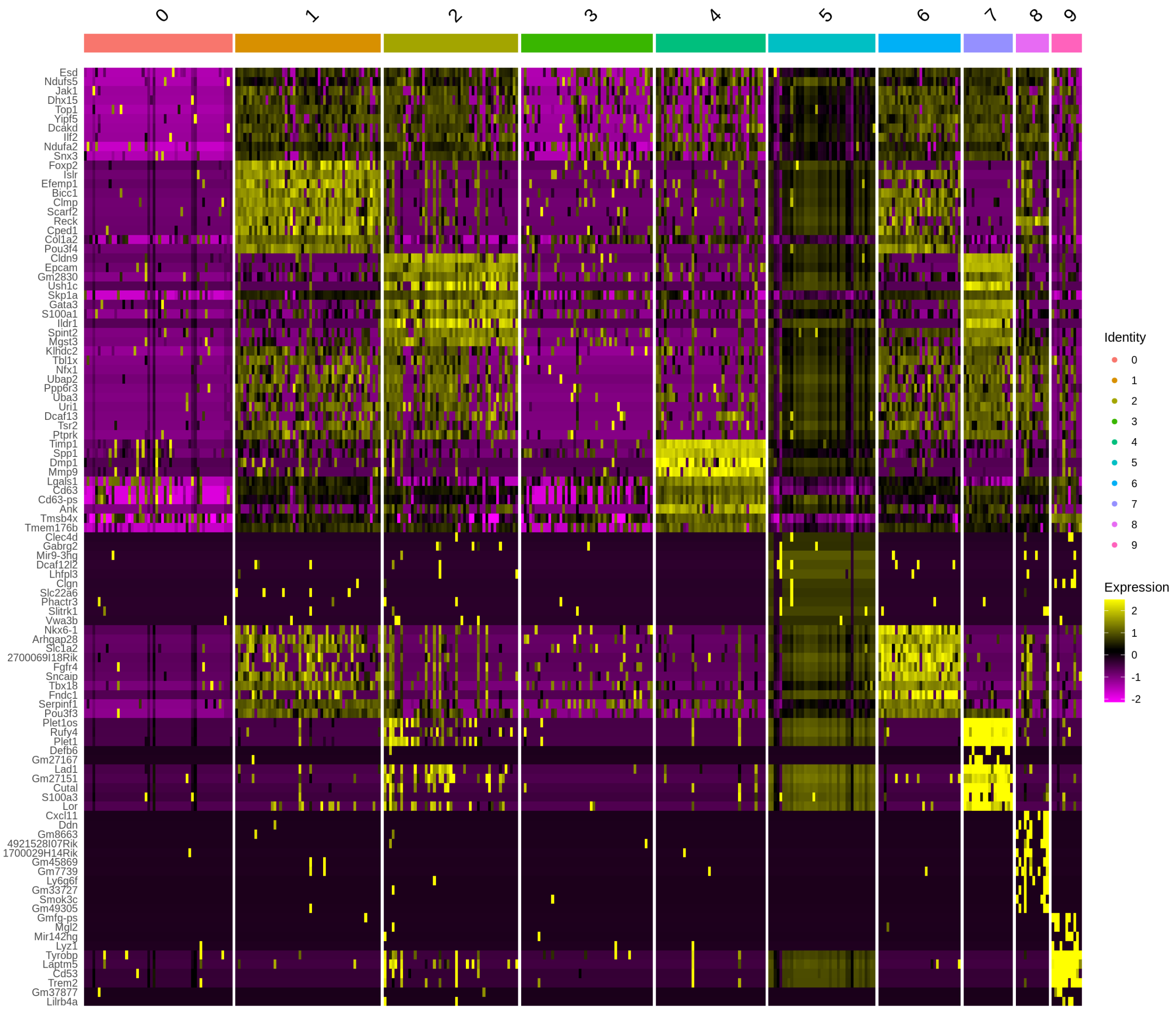

### S11 Fig

#### Cluster S1

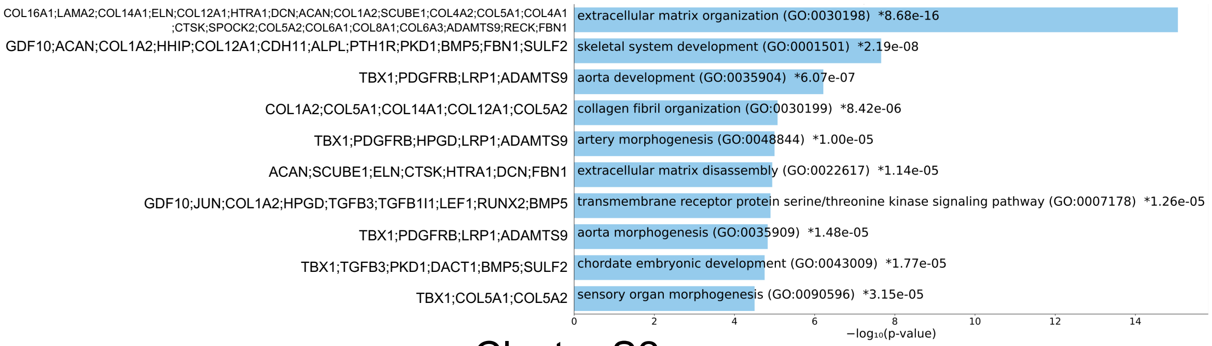


#### Cluster S2

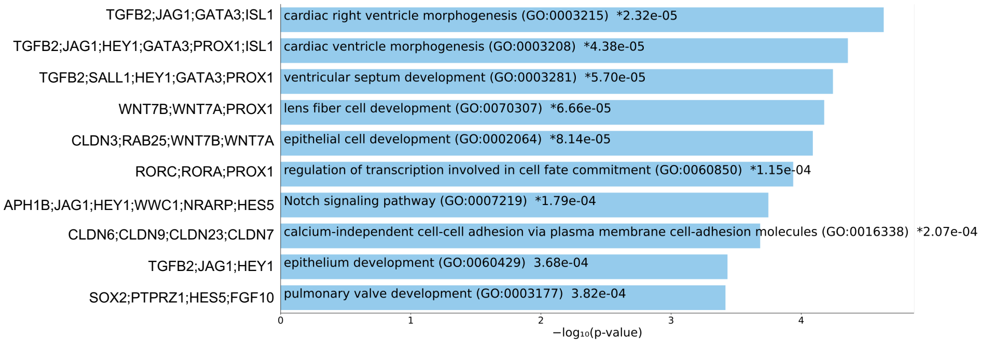


#### Cluster S6

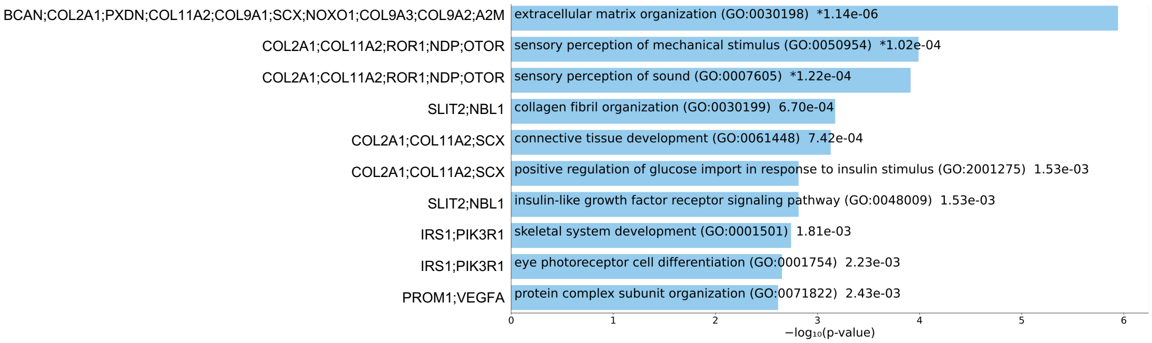


#### Cluster S8

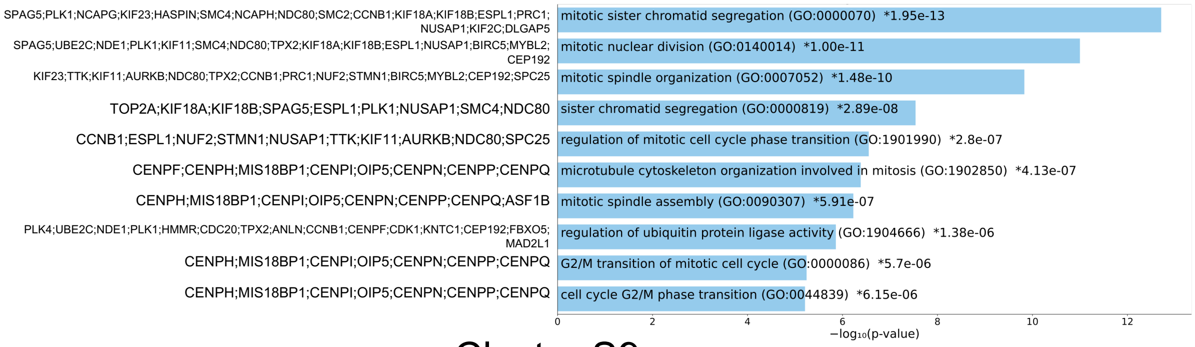


#### Cluster S9

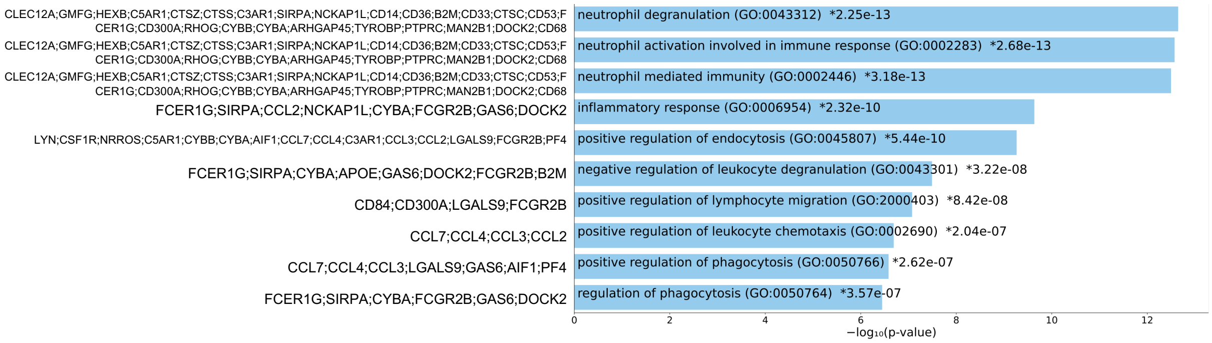

### S13 Fig

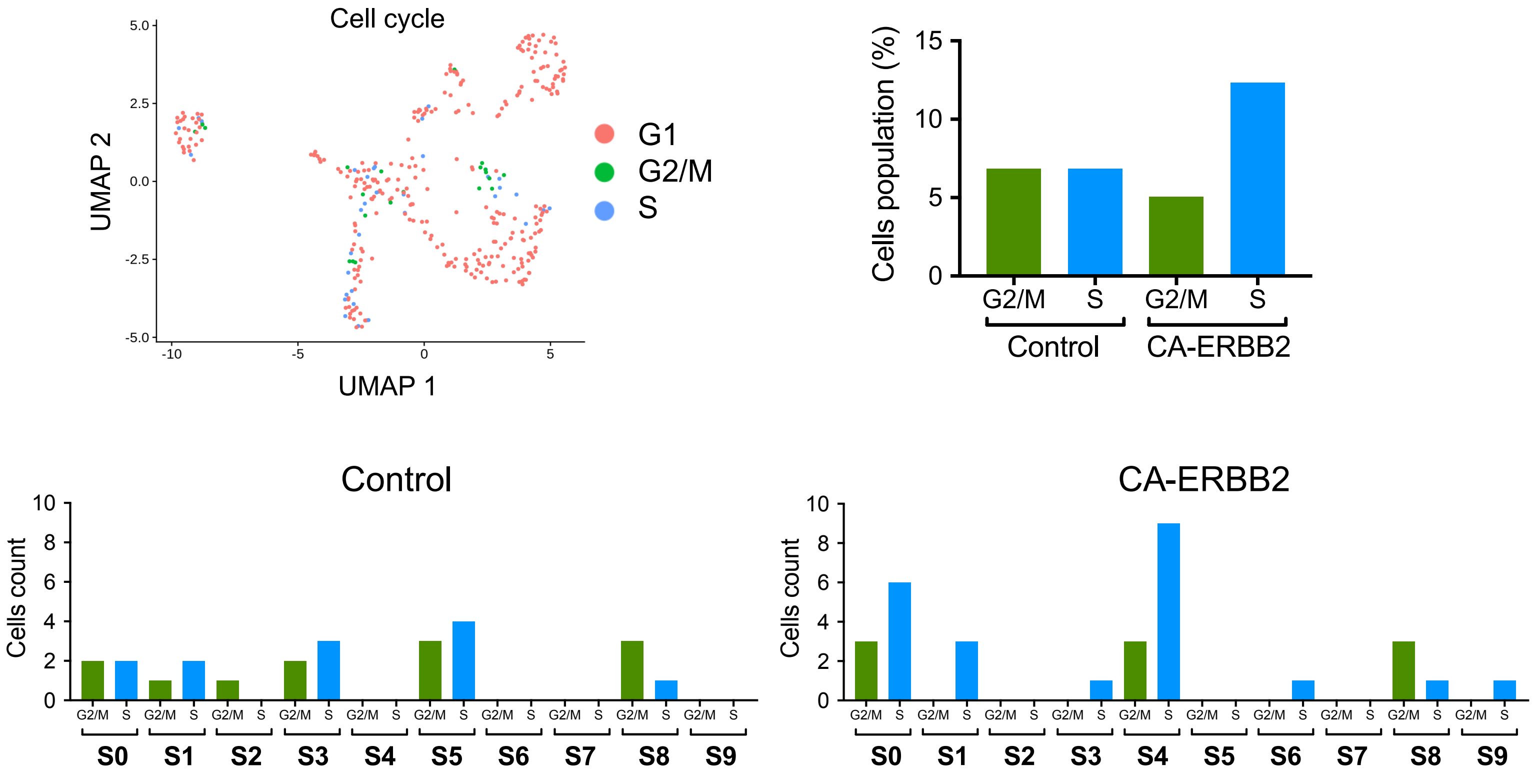
