## Supplementary material for "Single cell RNA sequencing analysis of mouse cochlear supporting cell transcriptomes with activated ERBB2 receptor, a candidate mediator of cochlear regeneration mechanisms": S4 Table

**S4 Table: Cell-type specific markers**. Gene markers were selected based on (Kolla et al, 2020).

| | | |
|---|---|---|
| DC1/2 | Dieter Cells from rows 1 and 2 | Hes5, Pdzk1ip1, Ppp1r2, S100a1, Serpine2 |
| DC3 | Dieter Cells from row 3 | Hes5, Igfpl1, Lfng, Prss23, S100a1 |
| Hensen | Hensen Cells | Egfl6, Fam159b, Fst, Nupr1, Pmch |
| IHC | Inner Hair Cells | Ccer2, Pcp4, Cib2, Pvalb, Acbd7 |
| IPC | Inner Pillar Cells | Cryab, Emid1, Igfbpl1, Npy, S100b |
| IPhC | Inner Phalangeal Cells | Anxa5, Fabp7, Gjb2, Matn4, Prss23 |
| IS | Inner Sulcus Cells | Igf1, Matn1, Meg3, Rgcc, Rm4sf1 |
| IdC | Interdental Cells | Cdkn1c, Fxyd6, Otoa, Ptgds, Smoc2 |
| LGER1* | Lateral Greater Epithelial Ridge Cells, group 1 | Dcn, Ddost, Pdia6, Rcn3, Sdf2l1 |
| LGER2* | Lateral Greater Epithelial Ridge Cells, group 2 | Cpxm2, Ctgf, Fkbp9, Kazald1, Tectb |
| LGER3* | Lateral Greater Epithelial Ridge Cells, group 3 | Cst3, Gjb6, Net1, Tectb, Tsen15 |
| MGER | Medial Greater Epithelial Ridge Cells | Calb1, Crabp1, Epyc, Itm2a, Stmn2 |
| OHC | Outer Hair Cells | Insm1 |
| OPC | Outer Pillar Cells | Fam159b, Ppp1r2, S100b, Serpine2, Smagp |
| OS | Outer Sulcus Cells | Apoe, Bmp4, Fbln2, Fst, Npnt |
| Oc90 | Cells expressing Oc90 | Cndp2, Oc90, Rnase1, Ttr, Vmo1 |
