## Supplementary material for "Single cell RNA sequencing analysis of mouse cochlear supporting cell transcriptomes with activated ERBB2 receptor, a candidate mediator of cochlear regeneration mechanisms": S5 Fig

**A**

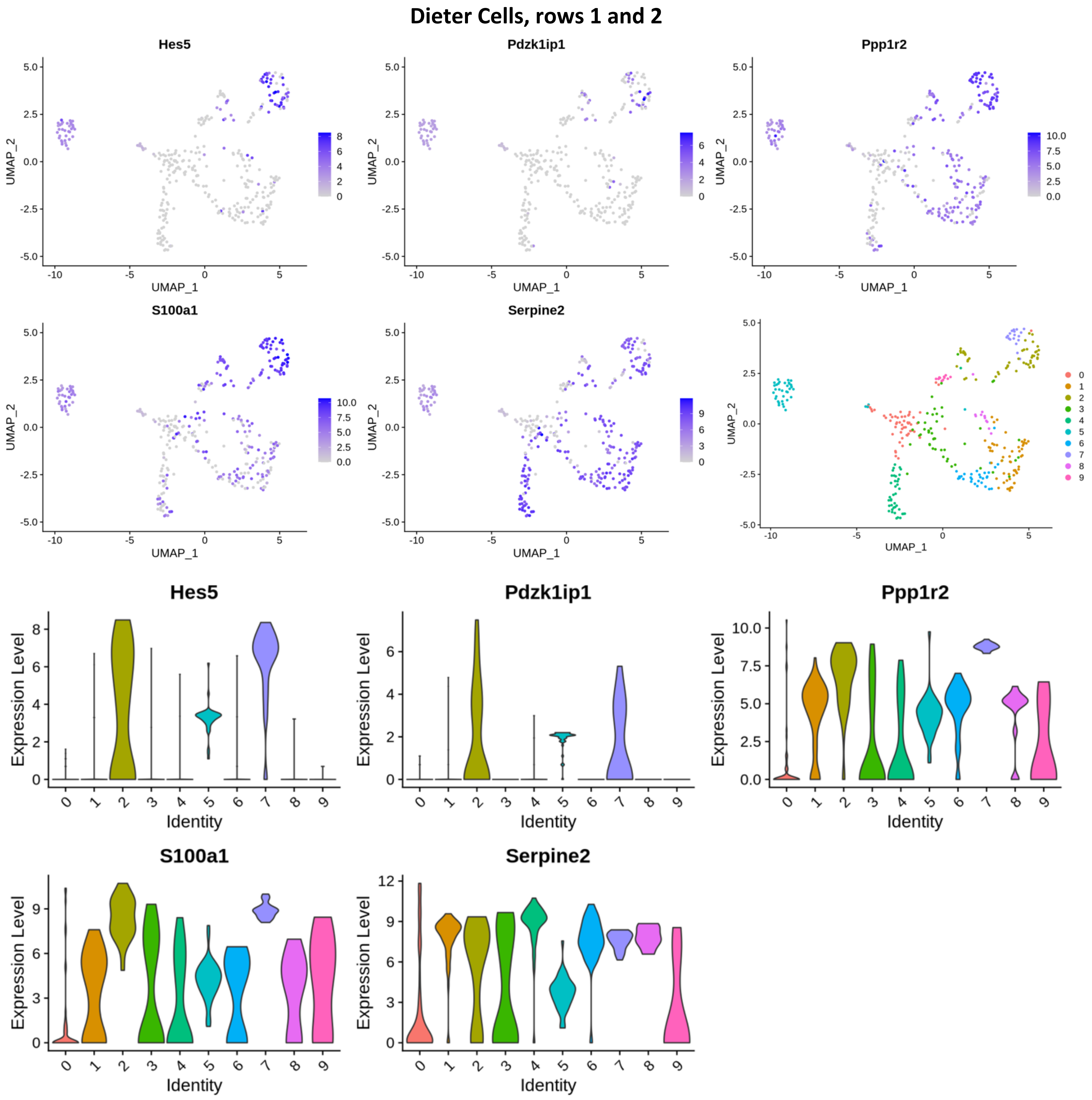

Dieter Cells, rows 1 and 2
Hes5
Pdzk1ip1
Ppp1r2
S100a1
Serpine2
UMAP_1
UMAP_2
Identity
Expression Level

**B**

### Dieter Cells, row 3

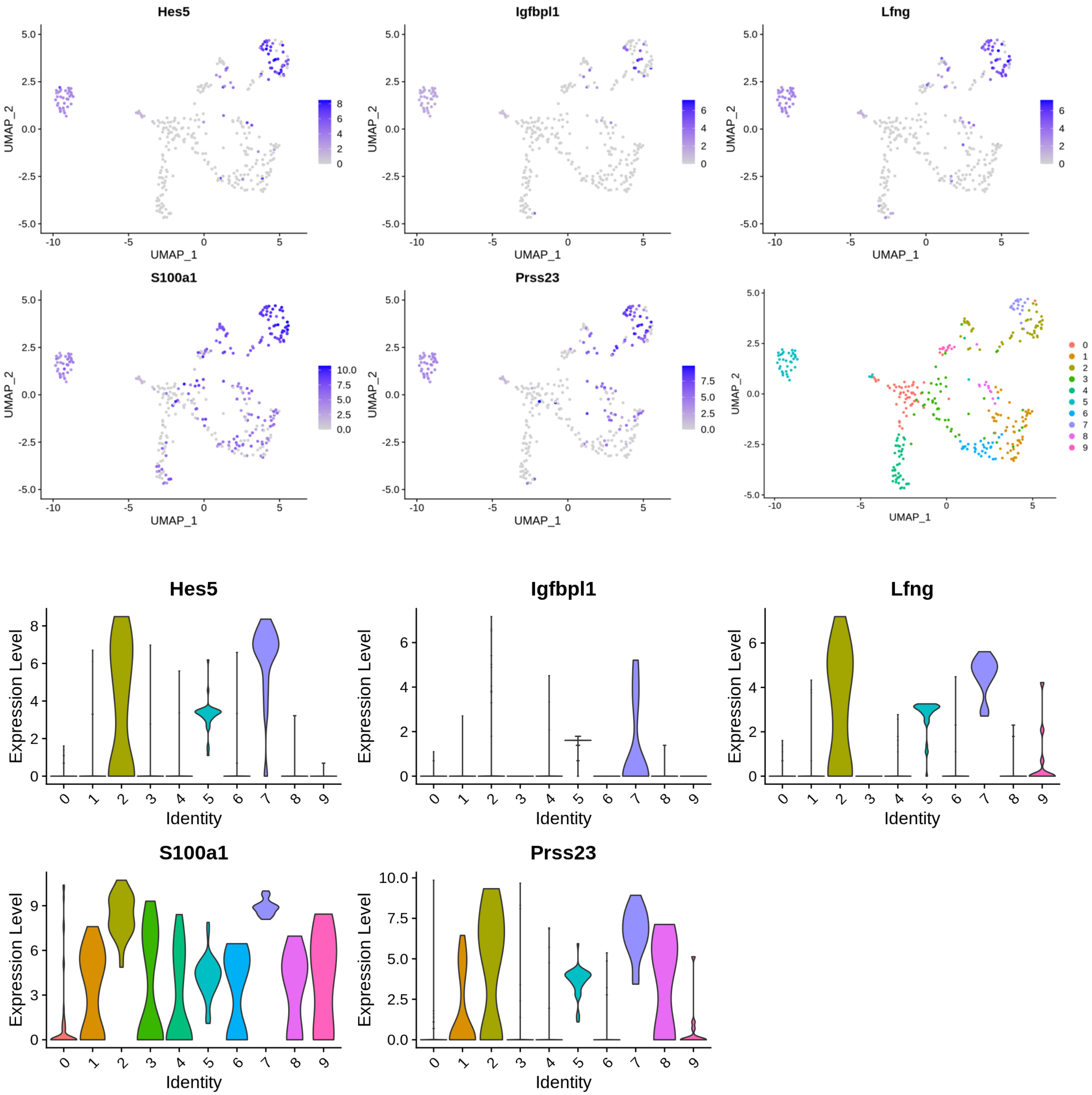

C

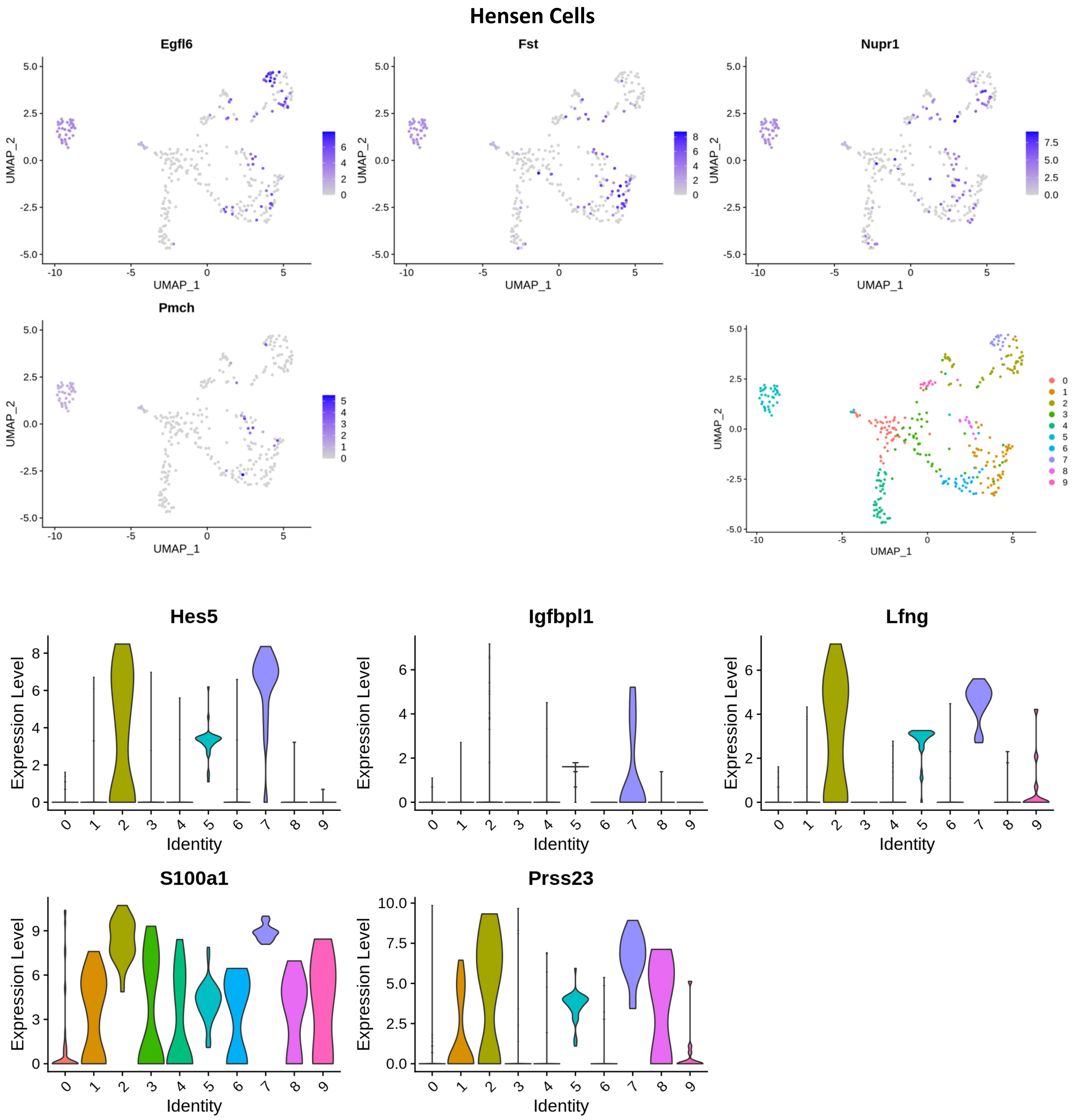
Hensen Cells
Egfl6
Fst
Nupr1
Pmch
UMAP_1
UMAP_2
Hes5
Igfbpl1
Lfng
S100a1
Prss23
Expression Level
Identity

**D**

### Inner Pillar Cells

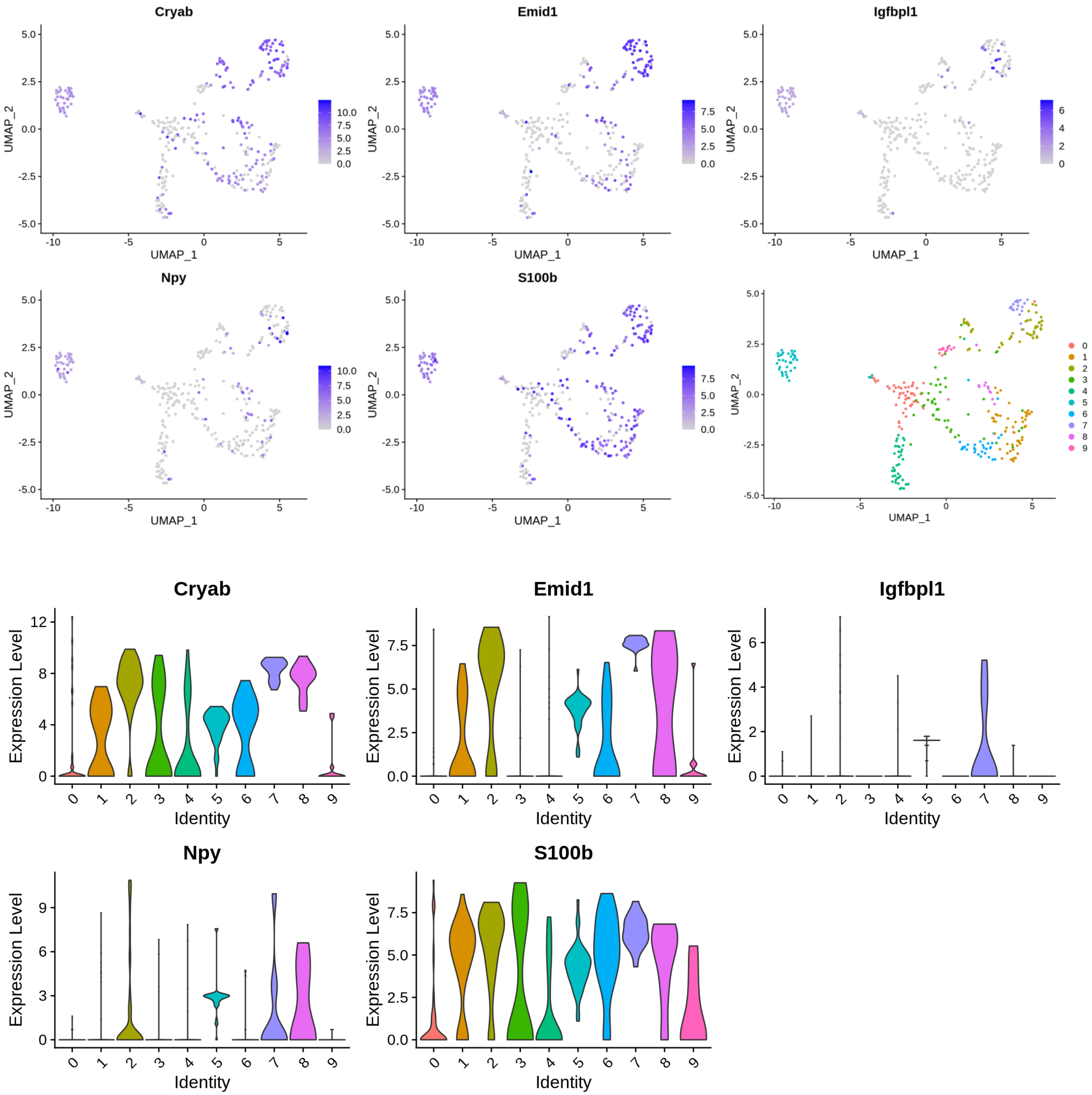

**E**

### Outer Pillar Cells

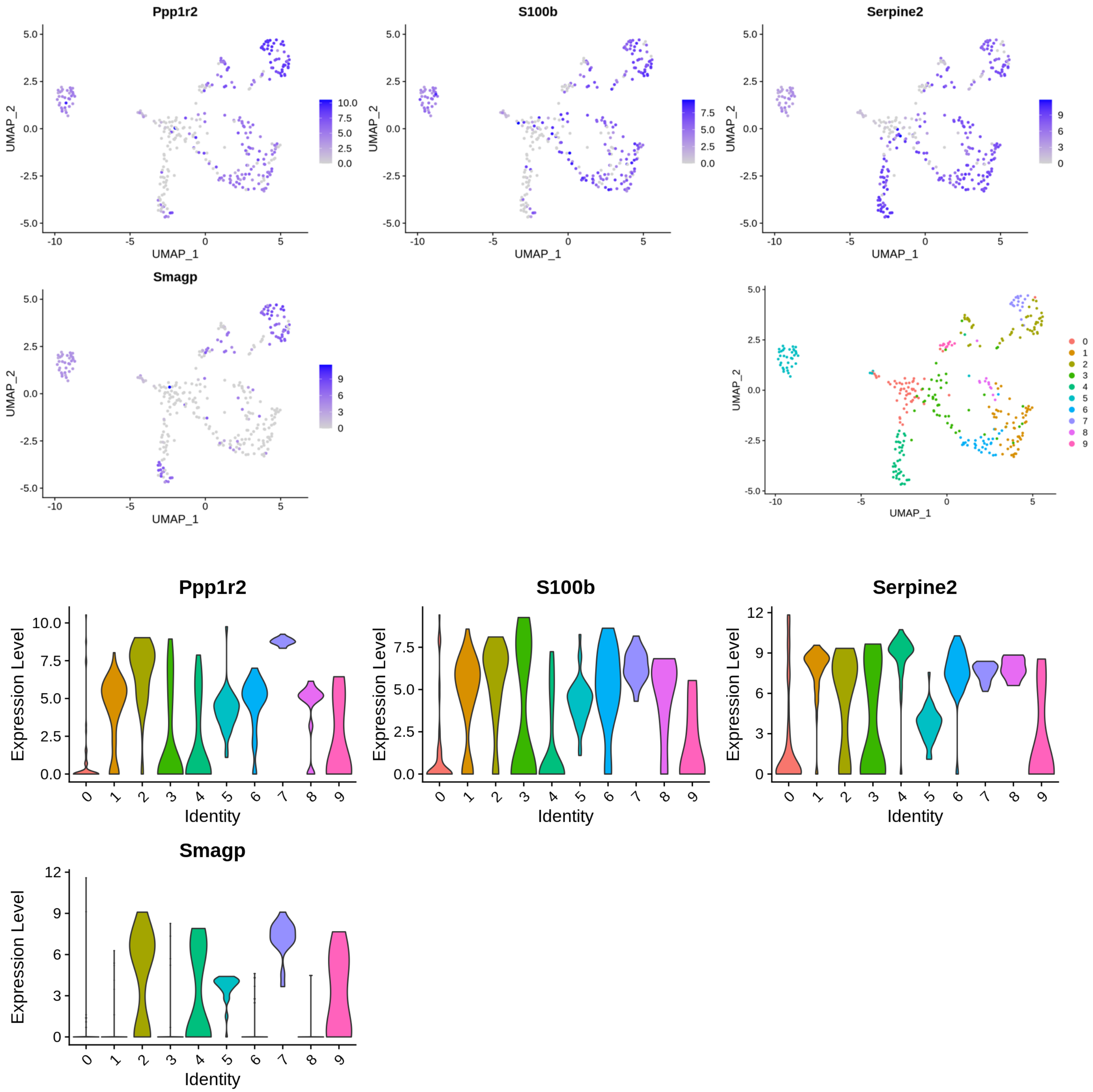

**F**

### Inner Hair Cells

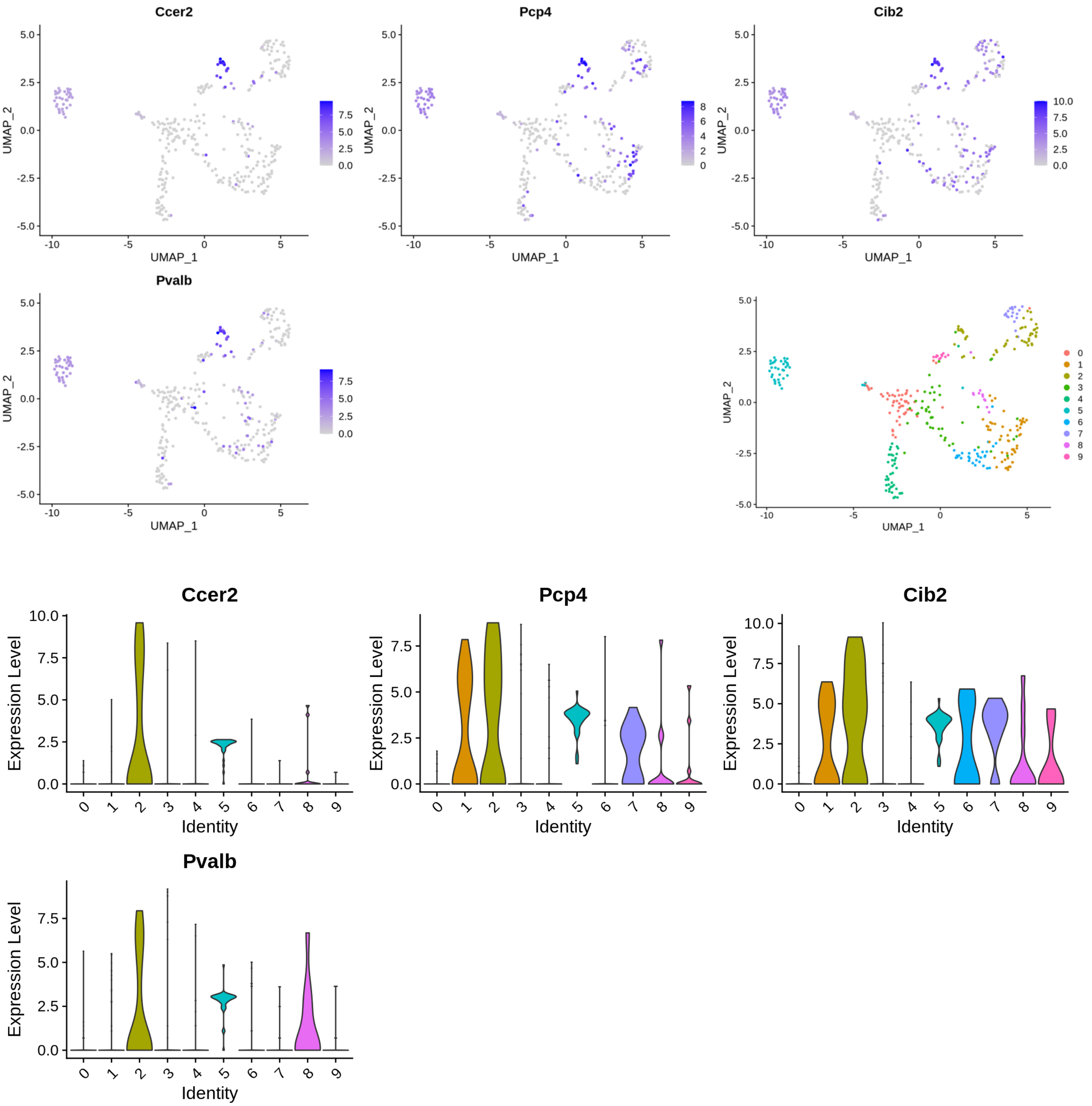

G

**Outer Hair Cells**

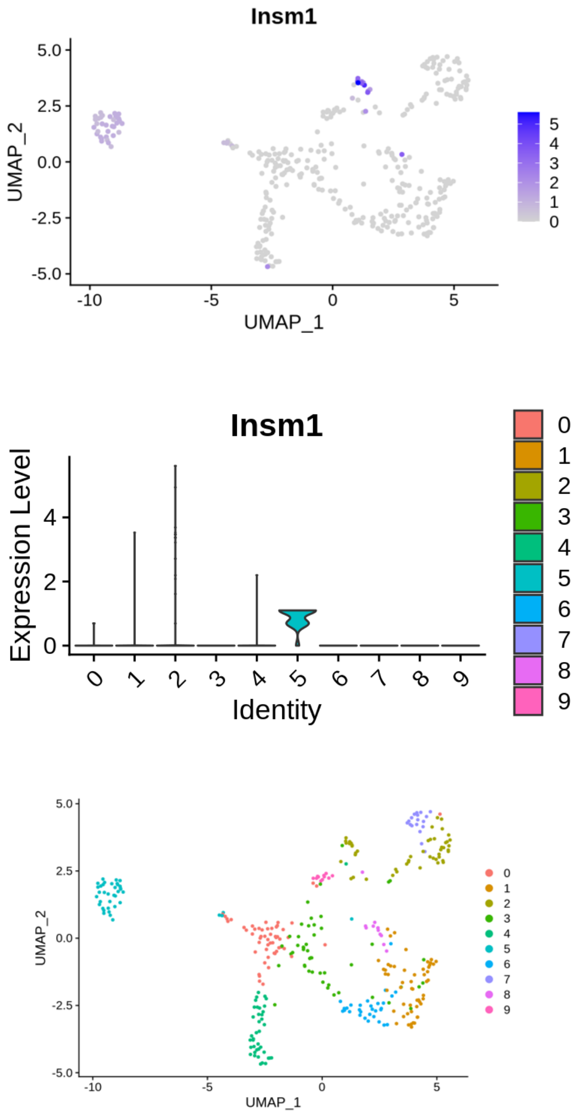

# H

### Lateral Greater Epithelial Ridge Cells, group 1

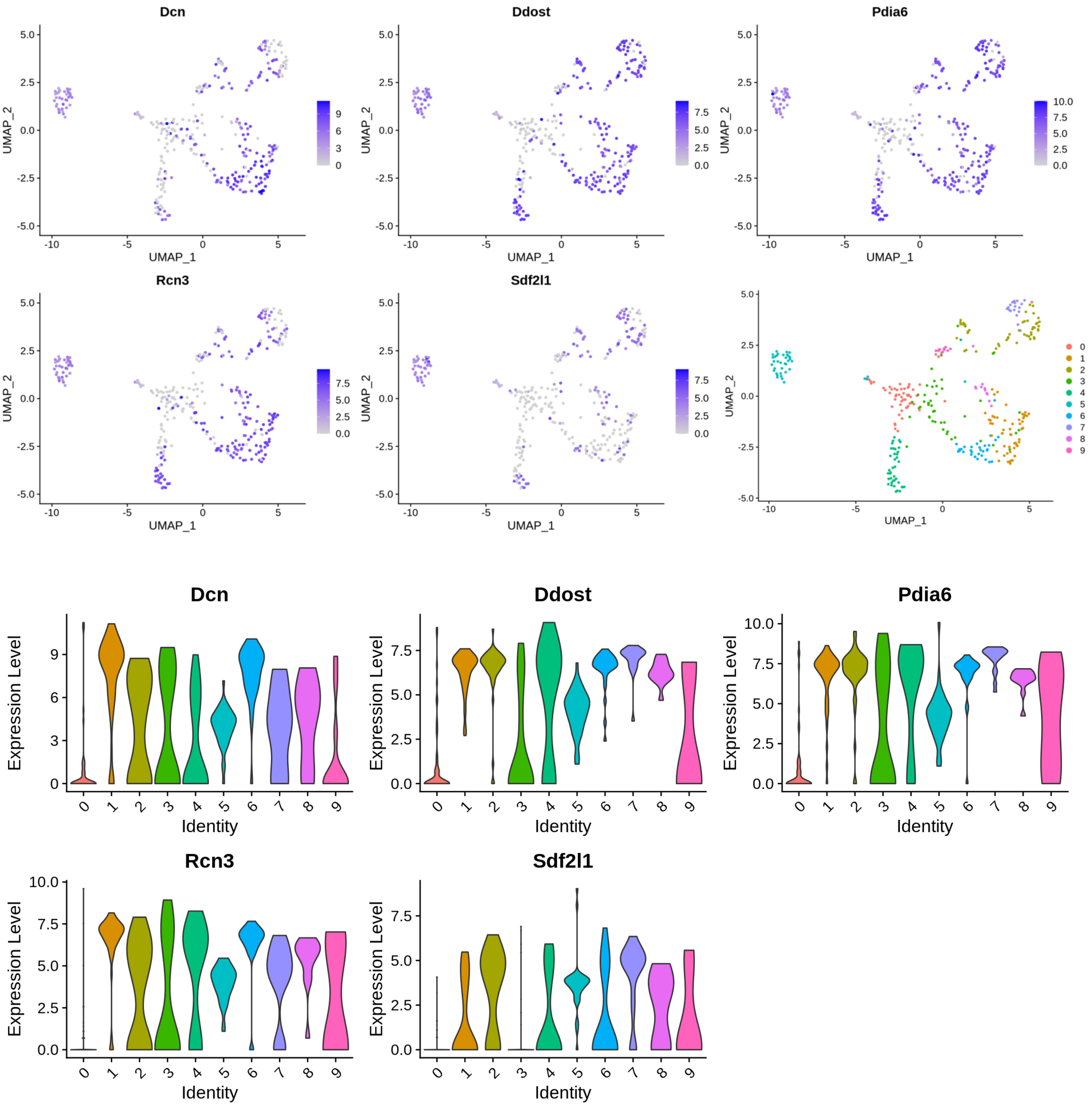

**I**

### Lateral Greater Epithelial Ridge Cells, group 2

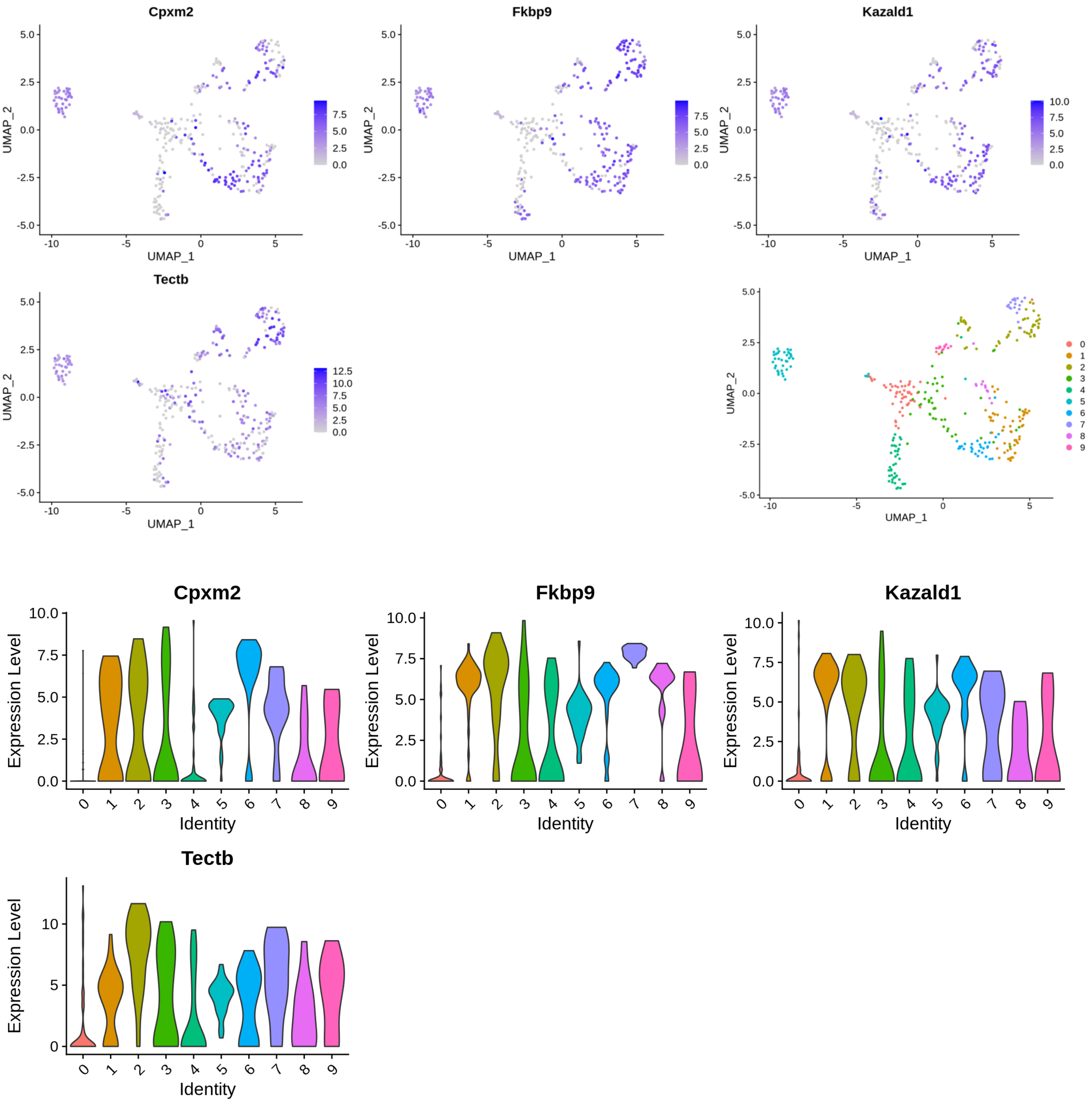

J

### Lateral Greater Epithelial Ridge Cells, group 3

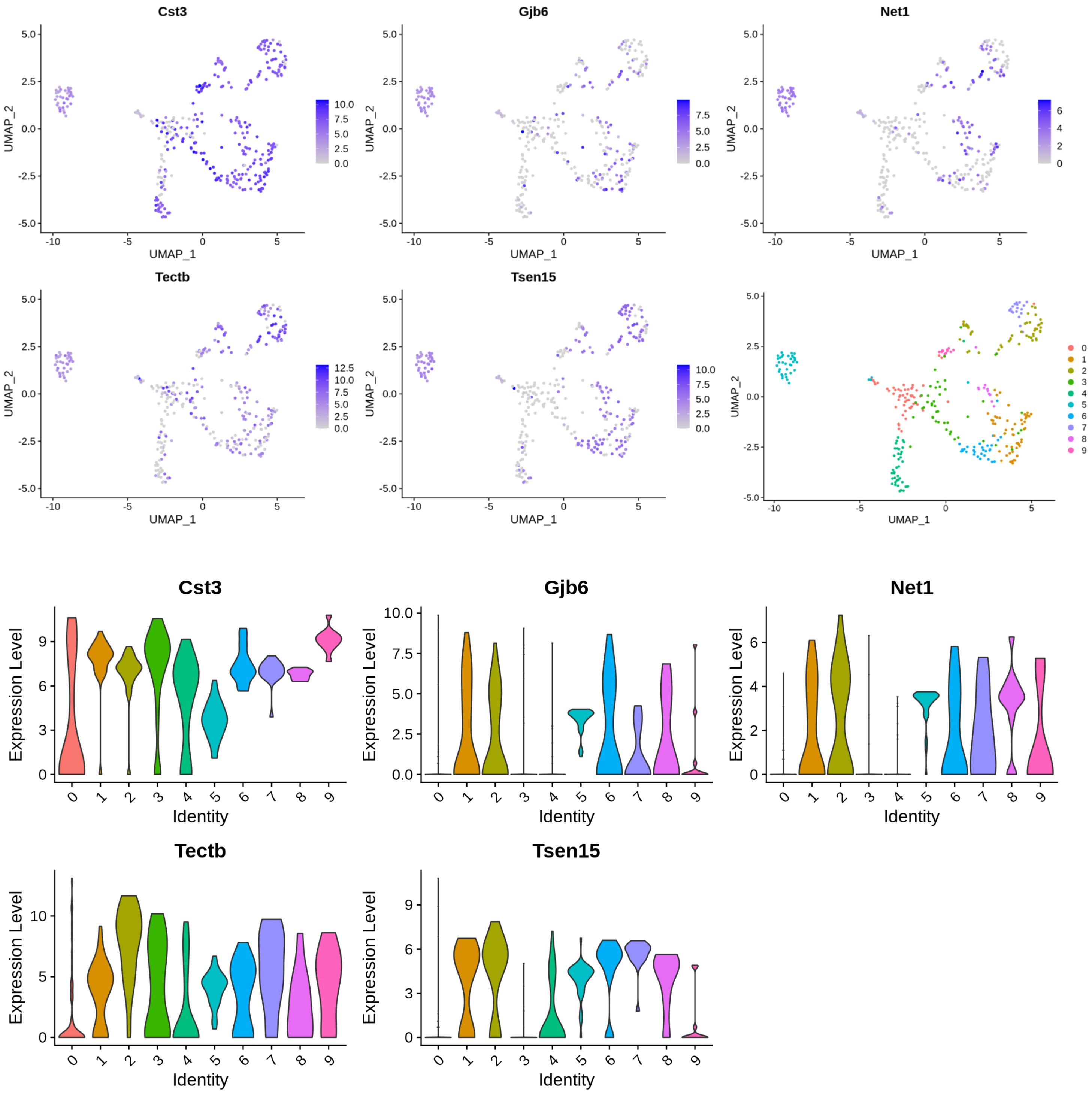

**K**

### Medial Greater Epithelial Ridge Cells

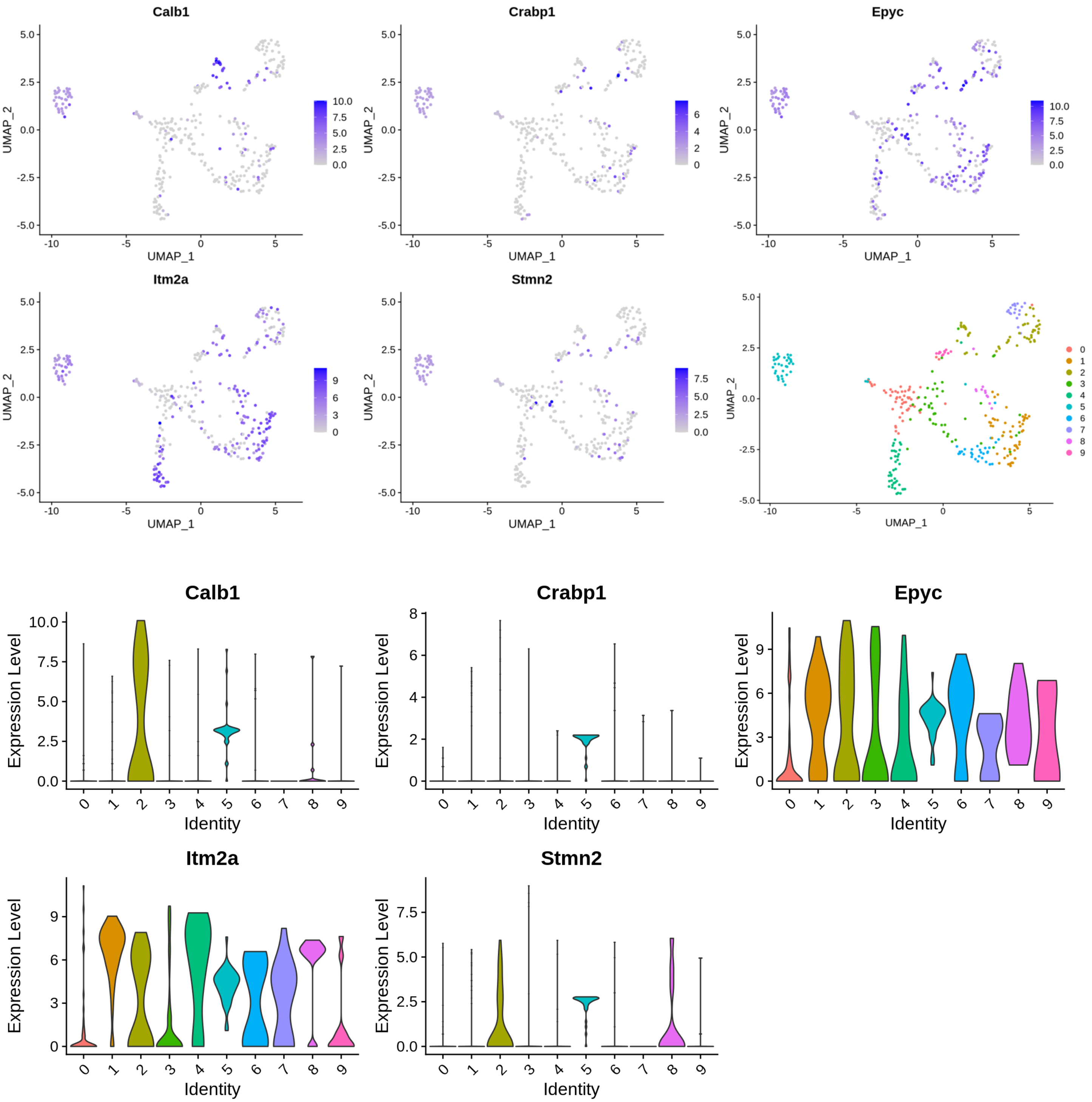

**L**

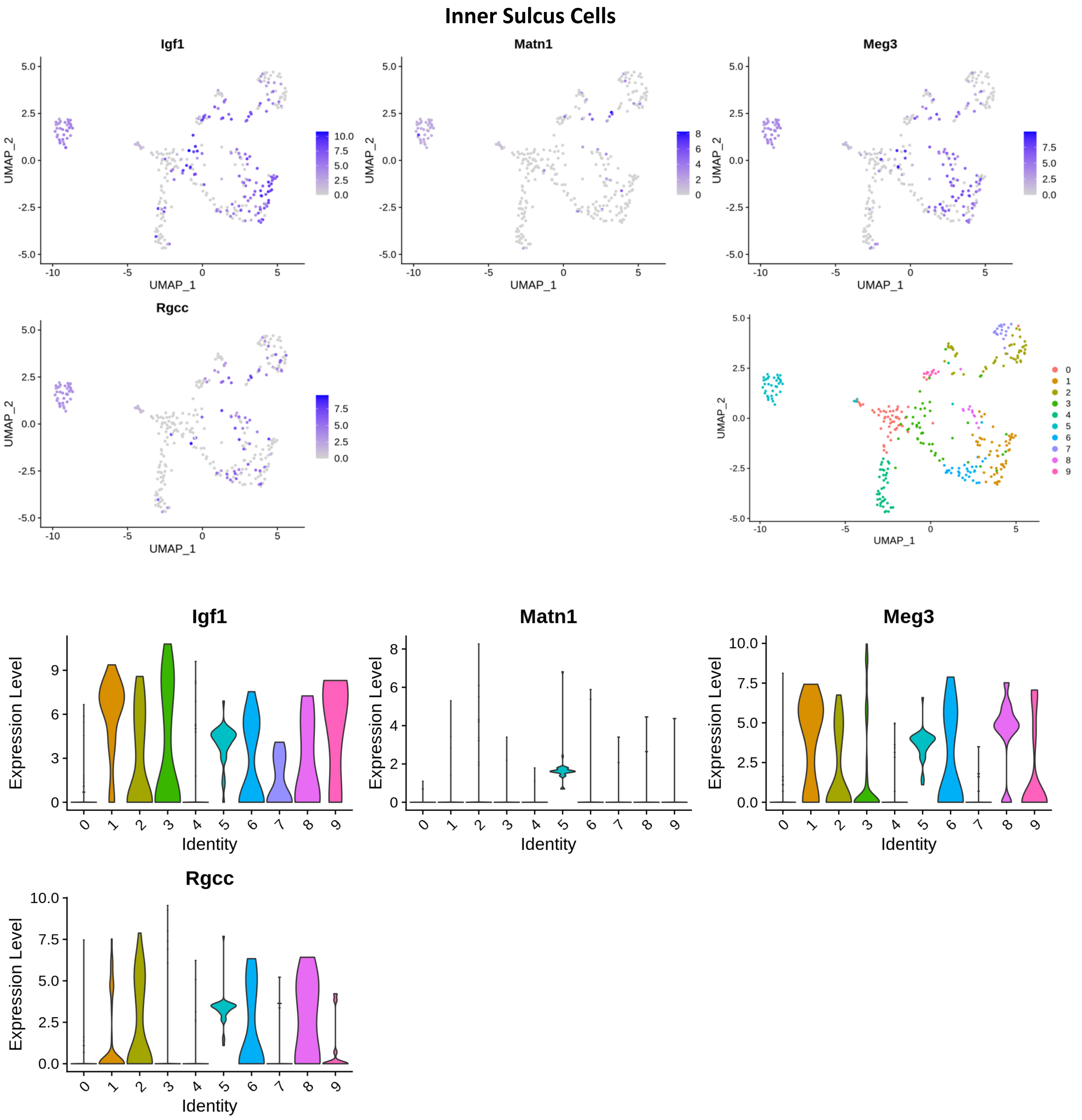
Inner Sulcus Cells
Igf1
Matn1
Meg3
Rgcc
UMAP_1
UMAP_2
Expression Level
Identity

**M**

### Outer Sulcus Cells

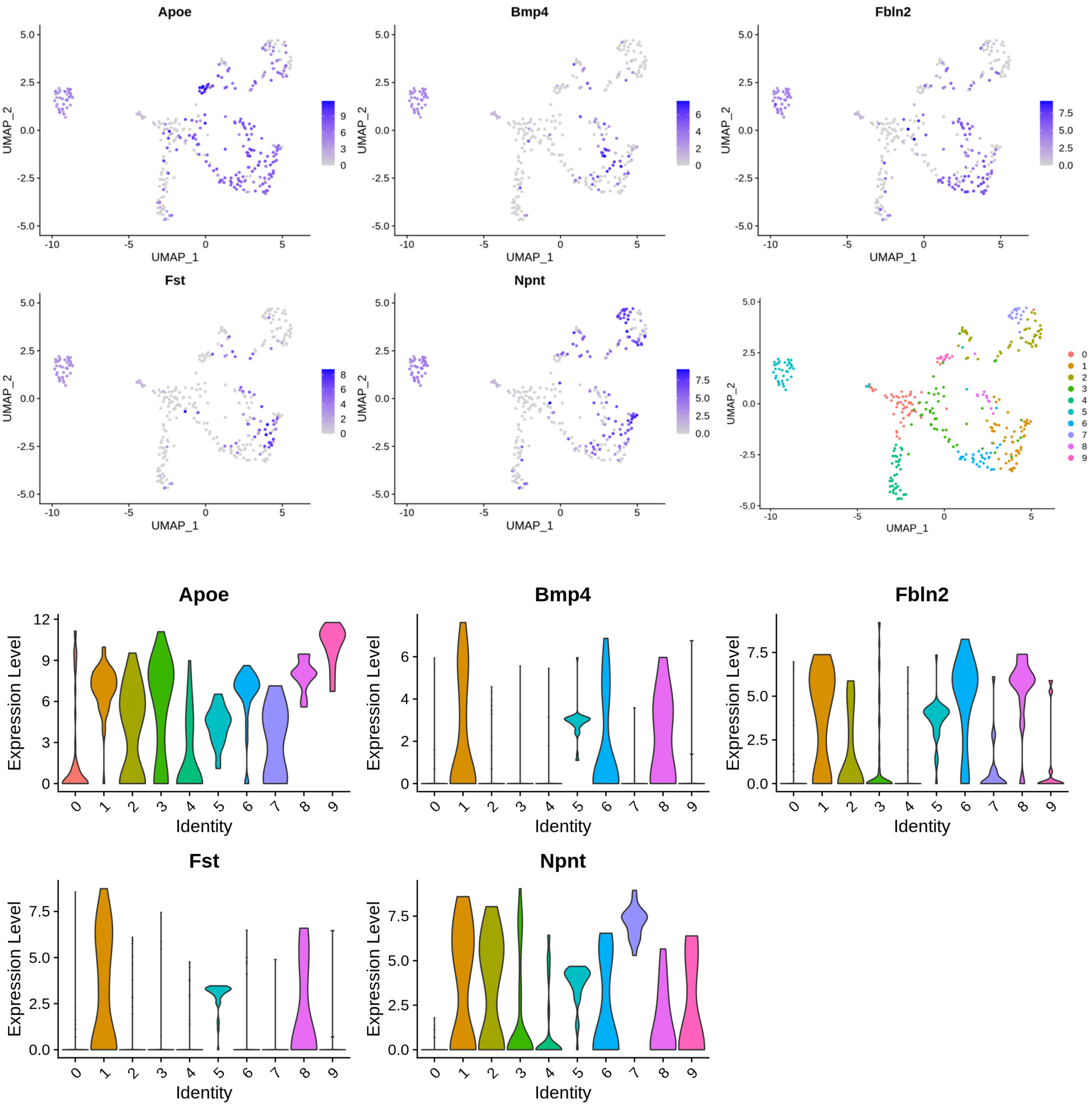

**N**

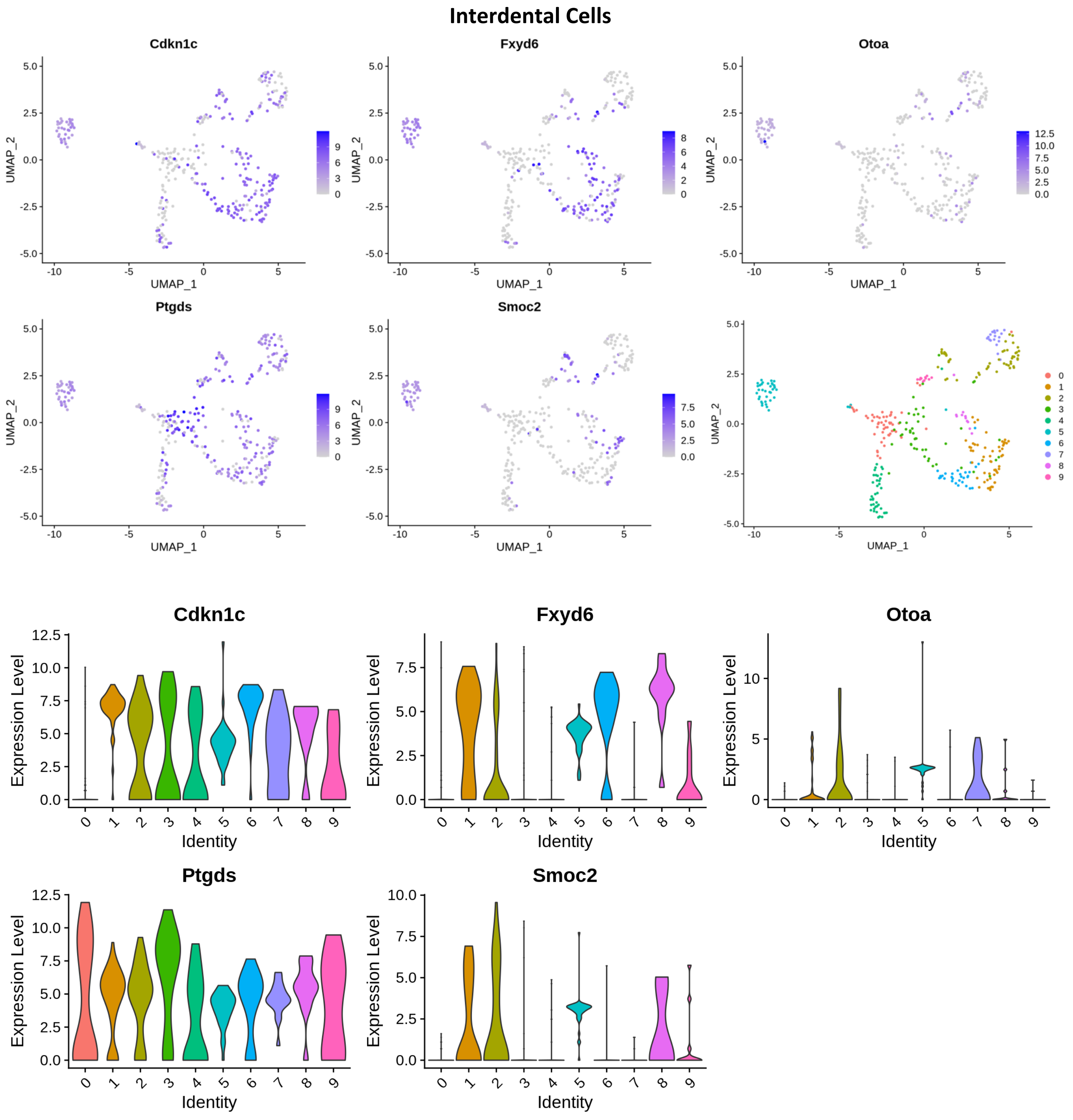
Interdental Cells
Cdkn1c
Fxyd6
Otoa
Ptgds
Smoc2
UMAP_1
UMAP_2
Expression Level
Identity

**O**

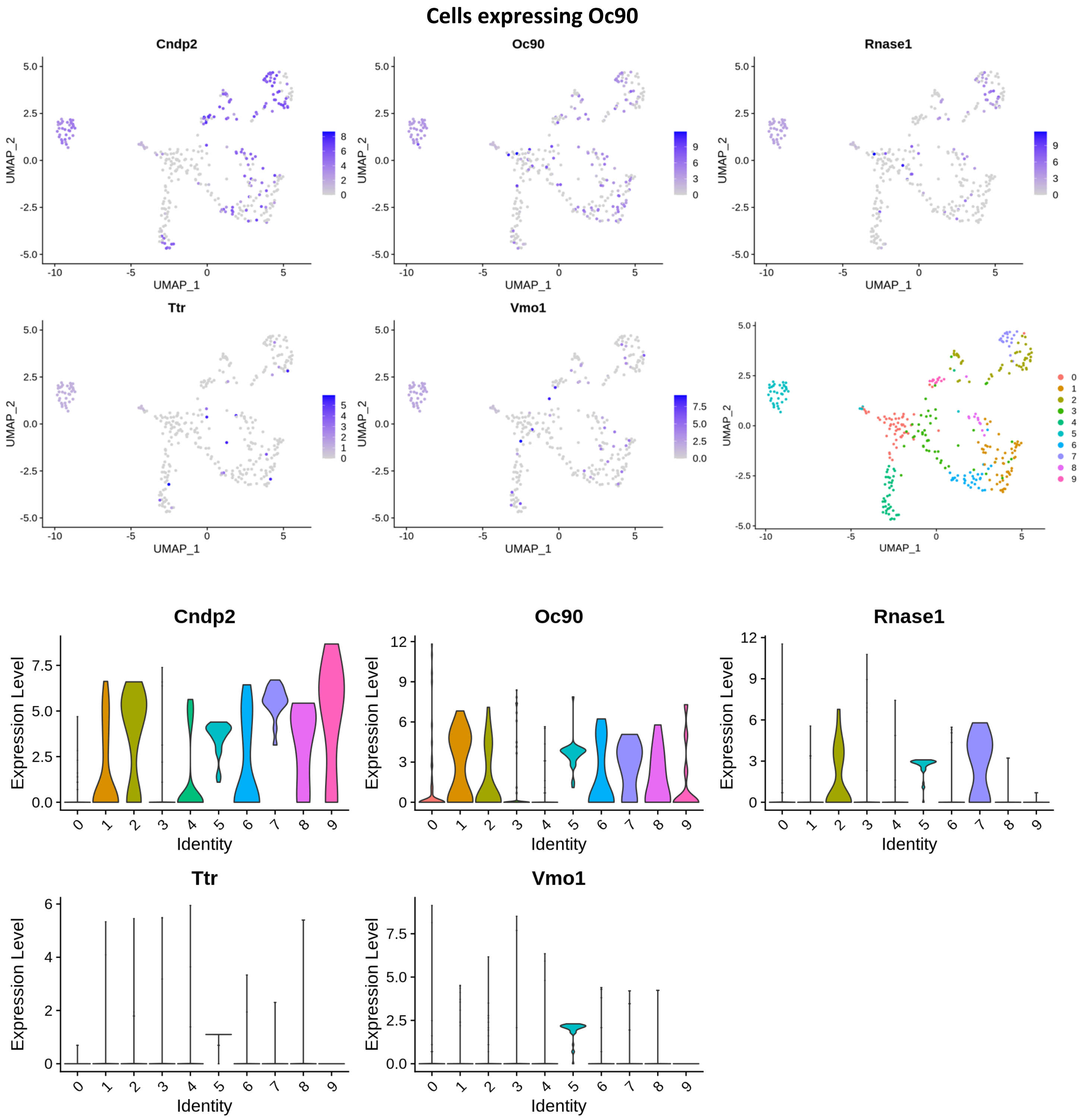
Cells expressing Oc90
Cndp2
Oc90
Rnase1
Ttr
Vmo1
UMAP_1
UMAP_2
Expression Level
Identity
