## Supplementary material for "Single cell RNA sequencing analysis of mouse cochlear supporting cell transcriptomes with activated ERBB2 receptor, a candidate mediator of cochlear regeneration mechanisms": S14 Fig

**S14 Fig. Additional validation of up-regulated genes.** RT-qPCR was performed on GFP+ FACS-sorted cochlear cells from P3 pups. The level of gene expression in CA-ERBB2 sample is presented as ΔΔCt value (± SD) relative to Control (CTRL) normalized against the expression of the two reference genes, *Eef1a1* (E) and *Tubb4a* (T) (n=3). Significance (**p*<0.05; ** *p*<0.01; *** *p*<0.001) was determined by unpaired *t*-test (one-tailed) analysis. Below, UMAP plots and split violin plots showing up-regulated expression of *Bglap, Bglap2, Ptgis, Il12a, CD63* and *Ank* in cells with activated CA-ERBB2.

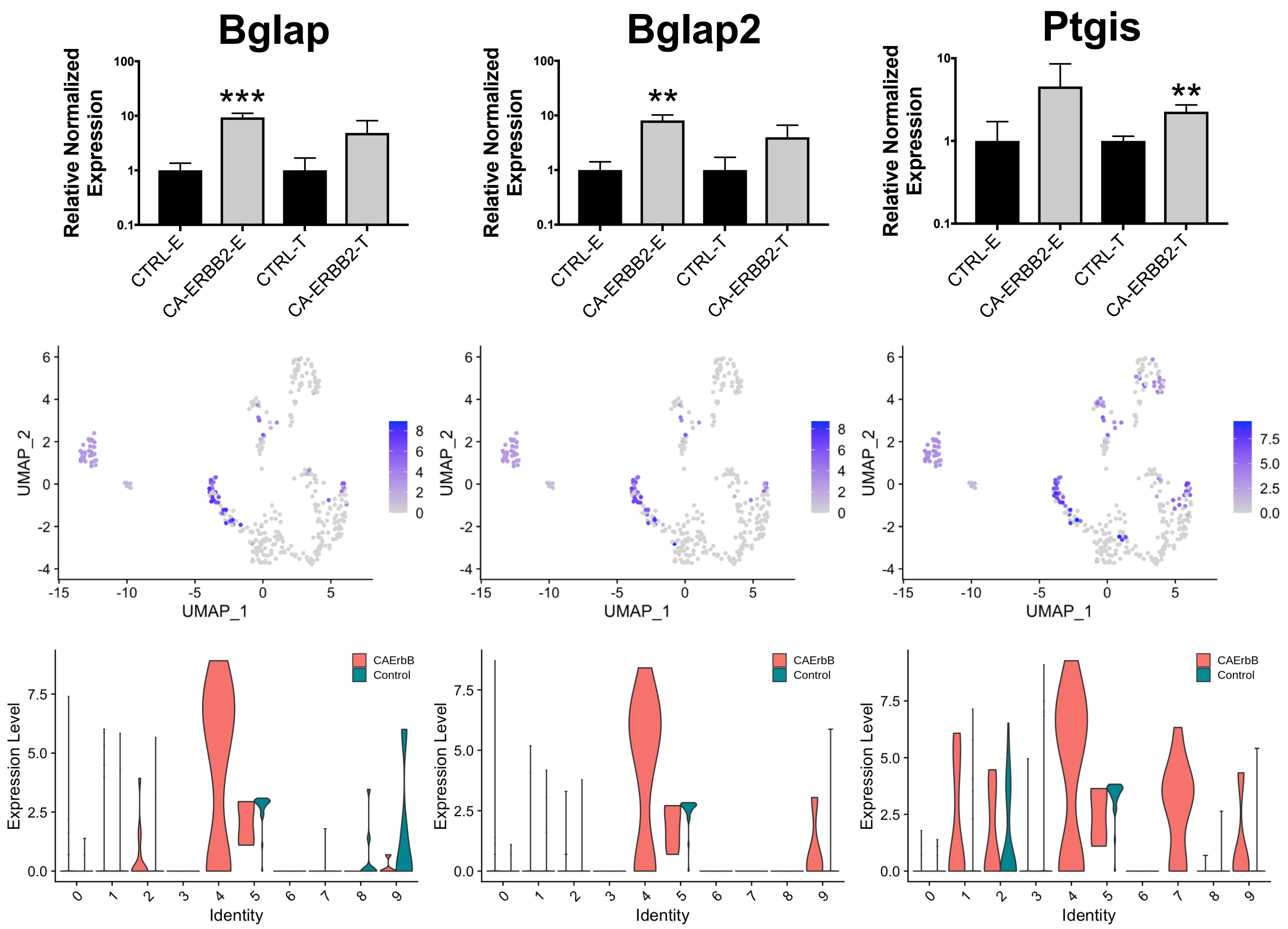

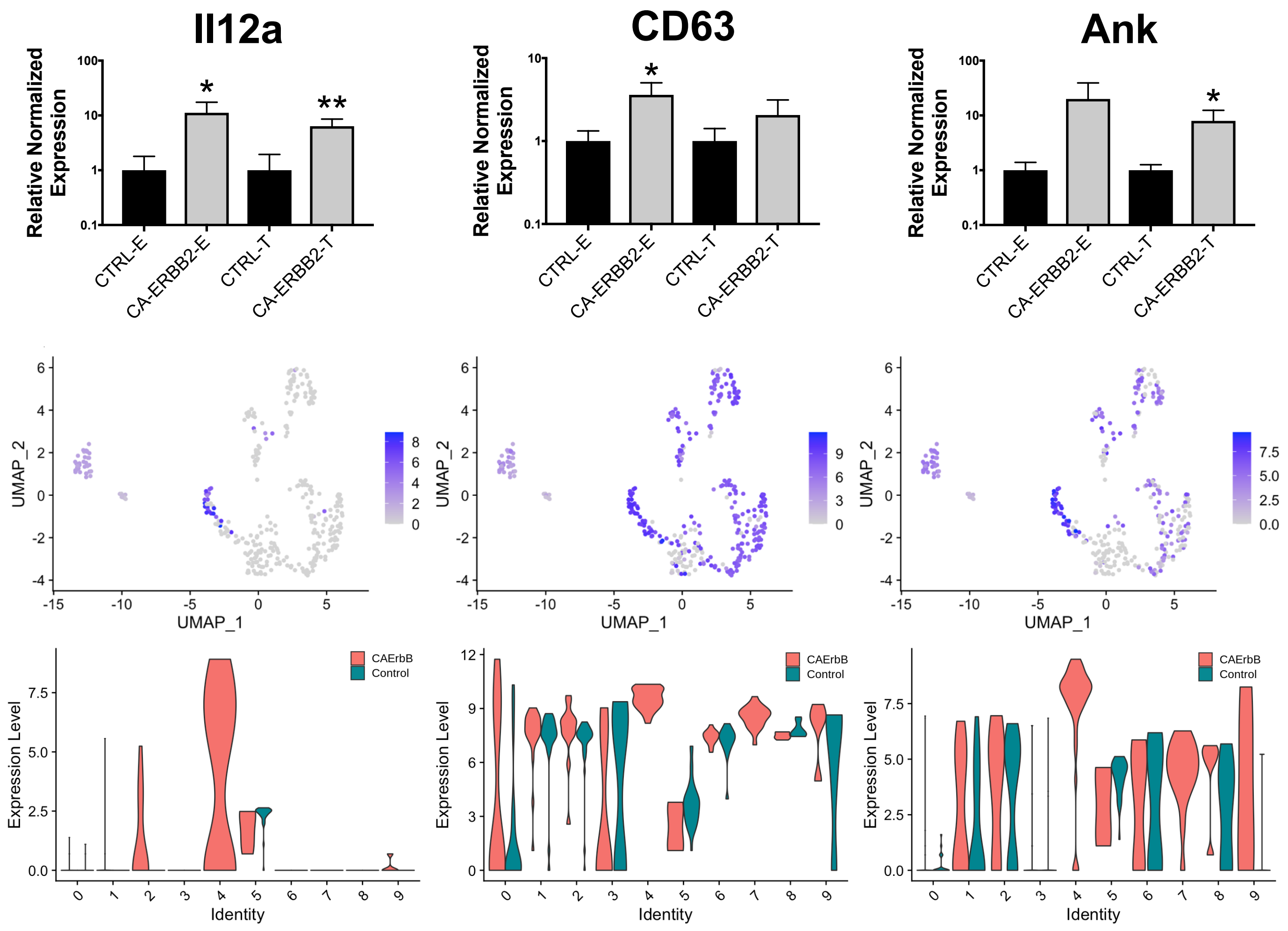
Il12a
CD63
Ank
Relative Normalized Expression
CTRL-E
CA-ERBB2-E
CTRL-T
CA-ERBB2-T
*
**
UMAP_1
UMAP_2
CAErbB
Control
Expression Level
Identity
