## Supplementary material for "Single cell RNA sequencing analysis of mouse cochlear supporting cell transcriptomes with activated ERBB2 receptor, a candidate mediator of cochlear regeneration mechanisms": S15 Fig

**S15 Table. Oligonucleotides used in genotyping and RT-qPCR analysis.**

| gene | Forward primer (5'-3') | Reverse primer (5'-3') |
|---|---|---|
| ***Genotyping*** | | |
| *CA-Erbb2* | AGCAGAGCTCGTTTAGTG | GGAGGCGGCGACATTGTC |
| *Fgfr3-iCre* | GAGGGACTACCTCCTGTACC | TGCCCAGAGTCATCCTTGGC |
| *Control* | CAAATGTTGCTTGTCTGGTG | GTCAGTCGAGTGCACAGTTT |
| ***RT-qPCR*** | | |
| *Tubb4a* | TGGACTCTGTTCGCTCAGGT | TGCCTCCTTCCGTACCACAT |
| *Eef1a1* | CAACATCGTCGTAATCGGACA | GTCTAAGACCCAGGCGTACTT |
| *Spp1* | GCTTGGCTTATGGACTGAGGTC | CCTTAGACTCACCGCTCTTCATG |
| *Dmp1* | CACGGACAGCAGTGAATCTGG | GCCGGTCCCCGTACTCTTA |
| *Mmp9* | GCTGACTACGATAAGGACGGCA | TAGTGGTGCAGGCAGAGTAGGA |
| *Timp1* | TCTTGGTTCCCTGGCGTACTCT | GTGAGTGTCACTCTCCAGTTTGC |
| *Bglap* | GCAATAAGGTAGTGAACAGACTCC | CCATAGATGCGTTTGTAGGCGG |
| *Bglap2* | GCAATAAGGTAGTGAACAGACTCC | GCGTTTGTAGGCGGTCTTCAAG |
| *Ptgis* | GGAGACAGGTCTCCTTGAGTTC | AACATCCGCTGAGTGGACACGA |
| *Il12a* | ACGAGAGTTGCCTGGCTACTAG | CCTCATAGATGCTACCAAGGCAC |
| *Cd63* | GGAATCCACTATCCATACCCAGG | CTCTTCACCAGACAGCAGGAGA |
| *Ank* | CGTGGACTCATGCTGGCATTCT | GTTCTCGGCATTCCAGGTGACT |
